# Evaluations of CRC2631 toxicity, tumor colonization, and genetic stability in the TRAMP prostate cancer model

**DOI:** 10.1101/2020.09.15.299081

**Authors:** Robert A. Kazmierczak, Bakul Dhagat-Mehta, Elke Gulden, Li Lee, Lixin Ma, Clintin P. Davis-Stober, Austen A. Barnett, Yves C. Chabu

## Abstract

Conventional cancer chemotherapies are not fully efficacious and do not target tumors, leading to significant treatment-related morbidities. A number of genetically attenuated cancer-targeting bacteria are being developed to safely target tumors *in vivo*. Here we report the toxicological, tumor-targeting, and efficacy profiles of *Salmonella enterica* serovar Typhimurium CRC2631 in a syngeneic and autochthonous TRAMP model of aggressive prostate cancer. CRC2631 preferentially colonize primary and metastatic tumors in the TRAMP animals. In addition, longitudinal whole genome sequencing studies of CRC2631 recovered from prostate tumor tissues demonstrate that CRC2631 is genetically stable. Moreover, tumor-targeted CRC2631 generates an anti-tumor immune response. Combination of CRC2631 with checkpoint blockade reduces metastasis burden. Collectively, these findings demonstrate a potential for CRC2631 in cancer immunotherapy strategies.

## Introduction

Conventional cancer chemotherapies are not specific and, as such, generate significant morbidities [1, 2]. Efforts to develop cancer-targeted therapeutics include the use of cancer-targeting bacteria to achieve cancer-specific cell killing. However, it has been a challenge to transition these bacteria-based approaches to the clinic due to a lack of a bacteria strain that is both safe and efficacious. Several bacterial strains have been developed, including the *Salmonella enterica* serovar Typhimurium strain VNP20009, one of the most studied tumor-targeting strains. VNP20009 was first isolated in a genetic screen for hyperinvasion mutants using a library of mutant strains derived from ultraviolet and chemical mutagenesis of strain 14028 [3]. Additional targeted genetic mutations were introduced in the *msb*, lipid A, and *purl* loci to attenuate VNP20009 and generate purine auxotrophy, respectively [4]. The safety and efficacy of VNP20009 were demonstrated in a wide range of pre-clinical animal cancer models [4-6], ultimately leading to clinical trials on metastatic melanoma or renal cell carcinoma patients [7]. The majority of VNP20009 pre-clinical studies relied on data derived from minimally aggressive tumors in immune-compromised animals [4-6], raising translatability concerns. Indeed, VNP20009 showed moderate toxicity but no anti-tumor effect in the aforementioned clinical studies [7], presumably because it was rapidly cleared by patients’ immune system. These studies have provided significant clinical insights and have underscored the need for cancer-targeting biologics that are not only safe and efficacious, but also likely to translate to the clinic.

We previously reported a tumor-targeting *Salmonella* typhimurium strain CRC2631 [8]. CRC2631 was derived from a parent strain (CRC1674) that was derived from the prototrophic wild-type *Salmonella* typhimurium LT2 strain [9] (Supplemental Table 1). CRC1674 was isolated in a genetic screen for mutants that selectively kill breast and prostate cancer cells in vitro using the Demerec collection [10]. This collection consists of mutant strains that arose naturally under nutrient-limiting conditions for over four decades, generating a wealth of genetically diverse and potentially attenuated strains [10-14]. CRC1674 was further attenuated by targeted deletion of *rfaH and thyA* genes. We also disrupted the *aroA* gene by Tn*10*d(Tc) transposon insertion. These modifications produced the attenuated strain CRC2631. *rfaH* is a positive transcriptional regulator of lipopolysaccharides (LPS) biosynthesis and its deletion [15, 16] lowers the expression of core LPS genes. The *aroA* transposon insertion and *thyA* deletion introduced auxotrophy for aromatic amino acids and thymine respectively [17-19].

Here, we report the toxicological and *in vivo* tumor-targeting profiles of CRC2631 in the syngeneic and autochthonous mouse model of aggressive prostate cancer, TRAMP (Transgenic Adenocarcinoma of Mouse Prostate). The B6FVB TRAMP model recapitulates some of the key genetic aspects of human prostate cancer. An androgen-dependent promoter drives the expression of *simian virus* 40 (SV40) large and small T antigens specifically in the mouse prostate epithelium. This leads to the inhibition of p53 and Rb, causing prostatic carcinomas by eight weeks of age. Similar to prostate cancers in men, these murine carcinomas disseminate throughout visceral organs, differentiate into neuroendocrine prostate cancer (NEPC), and ultimately kill the host [20-26]. While the molecular underpinnings that drive the conversion of carcinomas into NEPC are not well understood, NEPC is associated with loss of the tumor suppressors Rb and p53 in human prostate cancer [27].

In contrast to the B6 background, B6FVB TRAMP animals develop wide spread metastases [23, 28, 29], making them a suitable model to evaluate the therapeutic impact of CRC2631 on metastases.

We found that CRC2631 safely and persistently targets tumor lesions, including metastases. Longitudinal genome sequencing data from tumor-passaged CRC2631 revealed minimal genomic evolution. These findings indicate that CRC2631 is a genetically stable biologic that safely targets tumors. Moreover, tumor-targeted CRC2631 induces anti-tumor immune activity and concordantly reduces metastasis burden in the setting of checkpoint blockade.

## Results

### Evaluation of CRC2631 toxicity

VNP20009 is considered as the safety benchmark in bacterial cancer therapy development because it has been safely administered in human cancer patients [7, 30]. To determine the safety profile of CRC2631, we performed CRC2631 and VNP20009 comparative toxicological studies in TRAMP animals. We focused on treatment-related weight loss and lethality as toxicity measures. To control for tumor burden, groups of fourteen-week-old B6FVB TRAMP(+) mice were scanned by magnetic resonance imaging (MRI) and assigned either to the CRC2631 (*N=4*) or the VNP20009 (*N=4*) group. Animals received four weekly injections of 10^7^ CFU of CRC2631 or VNP20009 intraperitoneally (IP) (see methods, Supplemental Figure 1) and animal weight was monitored daily for four weeks. CRC2631 and VNP20009 had comparable effects on animal weight loss within the first two weeks of the study. During the last two weeks of the study, however, VNP20009-treated animals progressively lost more weight compared to CRC2631-treated animals (Figure 1a; *p*<0.0001, 3.50±1.77% versus 0.11±1.45% weight loss for VNP20009 and CRC2631, respectively). Consistent with CRC2631 being less toxic than VNP20009, the median survival time was 142 days for VNP20009 compared to 186 days for CRC2631 (Figure 1f). To more rigorously determine the toxicity of CRC2631 and derive its maximum tolerated dose (MTD), we escalated the dosing regimen to 10^7^ or 2.5×10^7^ or 5×10^7^ CFU administered every three days, instead of weekly, until 50% group lethality (LD50) was reached. B6FVB TRAMP(+) groups were treated IP (Figure 1b-c, 1e) or intravenously (IV) (Figure 1d) with a vehicle control (a sterile phosphate buffered saline, PBS) or 10^7^ or 2.5×10^7^ or 5×10^7^ CFU of CRC2631 or VNP20009. Compared to VNP20009, CRC2631 caused less weight loss across all dosage groups over the injection period. The average weight loss percentages for animals treated with 10^7^, 2.5×10^7^, and 5×10^7^ CFU CRC2631 were 7.20±2.45, 3.35±3.58, and 6.69±3.88, respectively. In contrast, VNP20009-treated animals exhibited an average weight loss of 8.99±2.56, 6.13±3.15, and 11.68±2.70 at the corresponding dose levels (Figure 1b, *p*<0.0085; Figure 1c, *p*<0.0001; Figure 1e, *p*<0.0139). Congruent with this, no lethality was observed in the CRC2631 group at the time the VNP20009-treated animals reached LD50 at the 10^7^ CFU/ three days dosage interval level (Figure 1g). In addition, the VNP20009 group experienced lethality after the first treatment at the 5×10^7^ CFU/ three days dose level, whereas the counterpart CRC2631 group exhibited no lethality until the third treatment when it precipitously reached LD50 (Figure 1h). This established the MTD at two doses of 5×10^7^ CFU administered three days apart.

**Figure 1.**
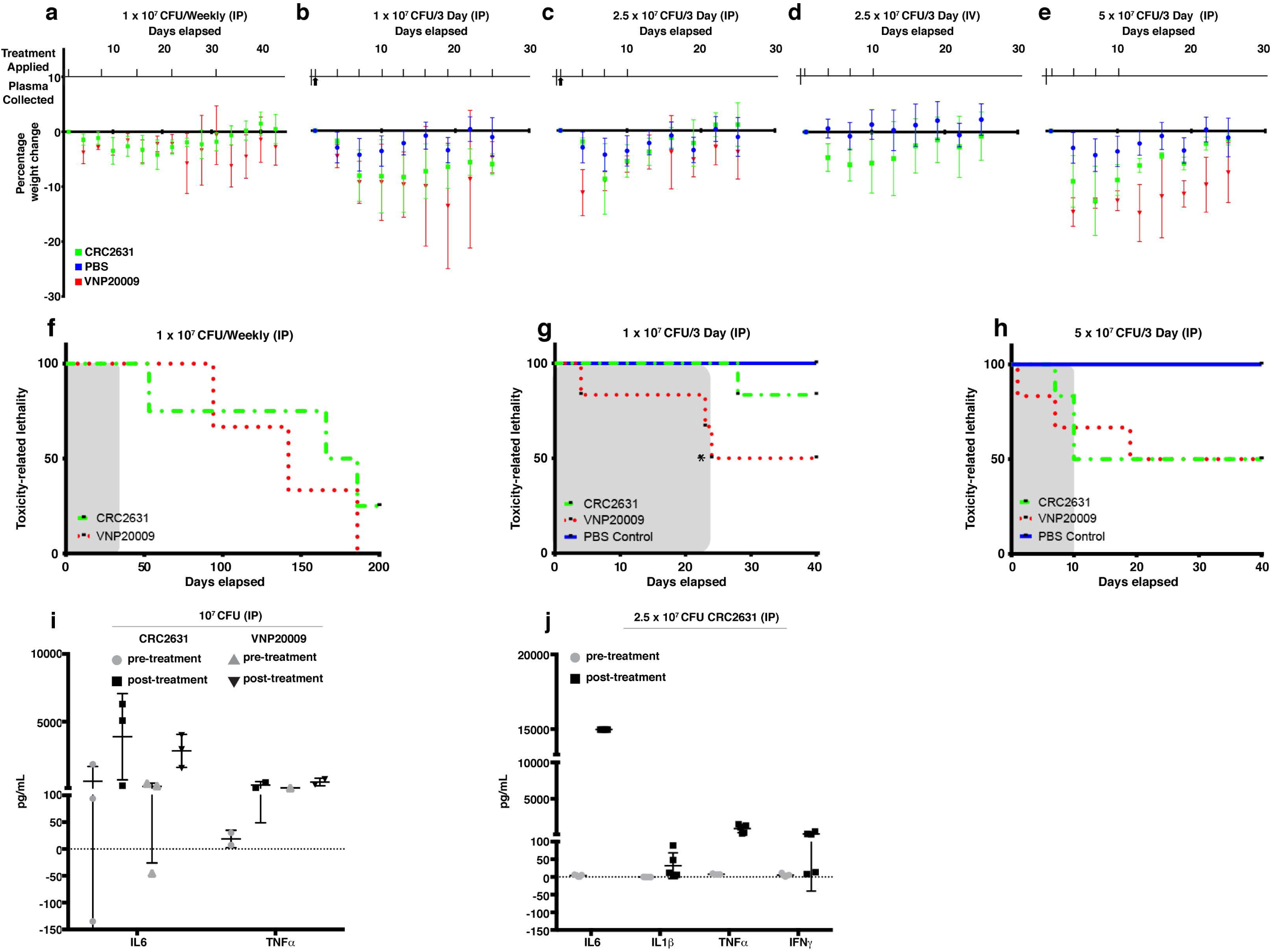
Comparative CRC2631 and VNP20009 toxicological assessment. Comparative toxicological assessment of CRC2631 and VNP20009 using mean group weight change, toxicity-related lethality, and cytokine response after treatment with CRC2631 or VNP20009 by intraperitoneal (IP) or intravenous (IV) injections into tumor-bearing B6FVB TRAMP(+) prostate cancer models. All B6FVB TRAMP(+) model tumor burdens were measured by MRI and mice groups sorted to normalize primary tumor volume size ranges for each dosage group before administration. Treatment application timing and sample harvesting schematics of CRC2631, VNP20009, or PBS vector control treatments during dosage frequency and concentration escalation experiments are shown for each experimental group in (a-e). Treatments applied at time points indicated by tick marks above x axis; whole blood samples for plasma extraction taken at time points indicated by upward arrows below x axis. (a) Mean percentage weight change of B6FVB TRAMP(+) mice (*N=4*) IP treated with 1×10^7^ CRC2631 (green) or VNP20009 (red) IP dosage every week for five weeks. (b, c, e) Mean percentage weight change of B6FVB TRAMP(+) mice (*N=6*) IP treated with (b) 1×10^7^, (c) 2.5×10^7^, or (e) 5×10^7^ CFU CRC2631 (green), VNP20009 (red), or equal volume PBS (blue) dosage every three days for 25 days or until LD50 was reached. Upward arrows indicate when plasma was sampled before and after treatment for profiling inflammatory cytokine response (i,j). (d) Mean percentage weight change of B6FVB TRAMP(+) mice (*N=12*) IV treated with four 2.5×10^7^ CFU CRC2631 (green) or equal volume PBS (blue) dosage every three days for 25 days. (f) Toxicological measure of survival over 200 days of B6FVB TRAMP(+) mice (*N=4*) IP treated with 1×10^7^ CRC2631 (green) or VNP20009 (red) every week for five weeks. Dosage time period shaded in gray. (g, h) Toxicological measures of survival over 40 days of B6FVB TRAMP(+) mice (*N*=6) IP treated with (g) 1×10^7^ or (h) 5×10^7^ CRC2631 (green), VNP20009 (red), or equal volume PBS (blue) dosage every three days for 42 days or until LD50 was reached (*compassionate euthanasia of mouse with >20% weight loss). Dosage time periods shaded in gray. (i) Inflammatory cytokine (IL-6, TNF*α*) immune response levels in B6FVB TRAMP(+) mice (*N=3*) plasma two hours before and two hours after first IP treatment using 1×10^7^ CRC2631 or VNP20009. (j) Inflammatory cytokine (IL-6, IL-1*β*, TNF*α*, INF*γ*) immune response levels in B6FVB TRAMP(+) mice (*N=5*) plasma two hours before and two hours after first IP treatment using 2.5×10^7^ CRC2631.

To minimize animal stress and thus the likelihood of animal lethality during the study, we performed all the remaining studies below MTD levels (i.e., at 10^7^ CFU or 2.5×10^7^ CFU per animal).

We asked whether the tolerability of CRC2631 is due to poor immunogenicity and it is not. CRC2631 and VNP20009 treatments triggered comparable cytokine responses in treatment naïve B6FVB TRAMP(+) mice plasma samples (Figure 1i, 1j).

Bacteria are cleared out of the blood circulation via the liver. VNP20009 causes significant liver toxicity in BALB/c mice bearing 4T1 mammary carcinoma xenografts. A single dose of 2×10^4^ CFU VNP20009 caused significant necrosis, inflammation, and extramedullary hematopoiesis (EMH) in liver tissue [31]. Thus, we sought to establish the impact of CRC2631 on liver pathology using a similar histopathological approach. Two groups of 31-week-old B6FVB TRAMP(+) mice (N=4) were treated IP with 4 doses of 2.5×10^7^ CRC2631 or 250ul PBS (control) at three-day intervals. Note that this dose is several orders of magnitude higher than what was used in the VNP20009 study. Liver histological sections were stained with hematoxylin and eosin (H&E) and examined by a veterinary pathologist. We detected no significant differences in necrosis, inflammation, and EMH between controls and CRC2631-treated animals (Figure 2b). Thus, in contrast to VNP20009, CRC2631 does not cause overt liver pathology.

**Figure 2.**
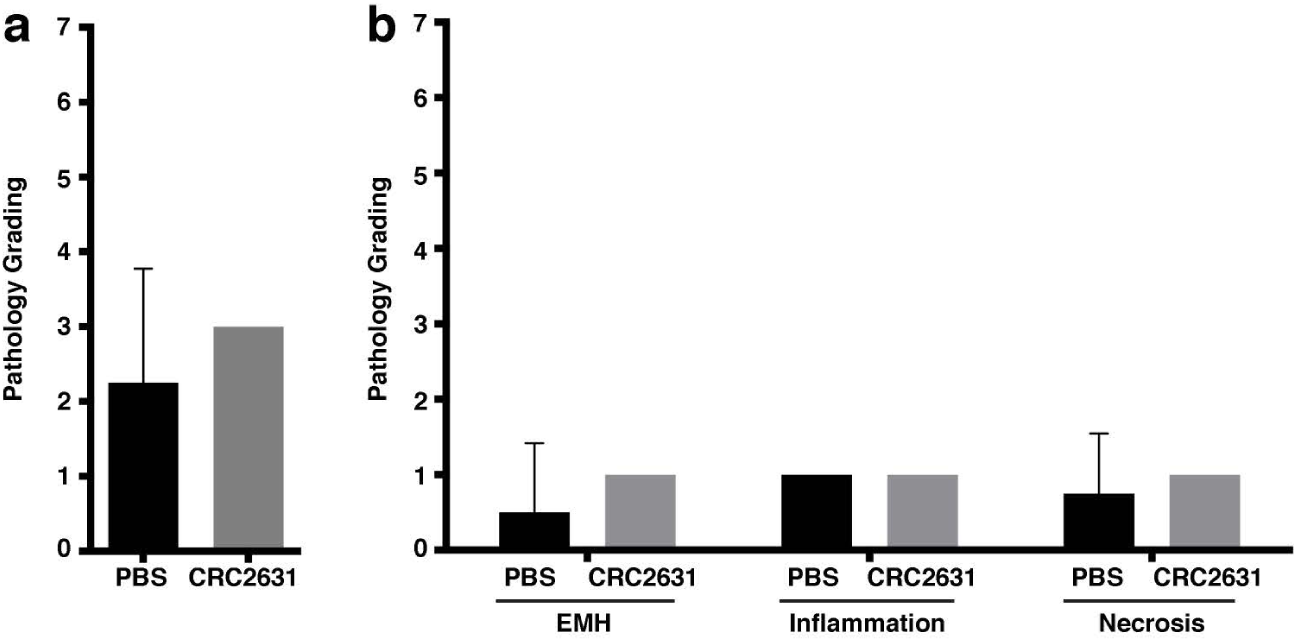
Histopathological analysis of non-targeted liver tissues in CRC2631 administered B6FVB TRAMP(+) mice. Histopathological analysis performed on hematoxylin- and eosin- (H&E) stained liver tissue samples obtained at necropsy from groups (*N=4*) of 33-week-old B6FVB TRAMP(+) mice. Groups were IP treated with no therapy control (PBS) or four doses of 2.5×10^7^ CRC2631 every three days (CRC2631) immediately followed by collection, mounting, and H&E staining of liver tissues to examine the effects of CRC2631 administration on non-targeted tissues. Pathology was graded based on levels of extramedullary hematopoiesis (EMH), observed amounts of inflammation, and necrosis in liver tissue. Pathology scoring key: 0 = no inflammation, EMH, or necrosis identified; 1 = up to 33% of the examined section had inflammation, EMH, or necrosis, 2 = 34%–66% of the examined section had inflammation, EMH, or necrosis; 3 = greater than 67% of the examined section had inflammation, EMH, or necrosis. Individual scores of EMH, inflammation, and necrosis were then summed to give a total composite score for each animal. (a) Mean composite liver tissue pathology score and standard deviation comparing PBS (no therapy) and CRC2631 (experimental) groups. (b) Liver tissue pathology scoring of the PBS and CRC2631-administered B6FVB TRAMP(+) groups examining EMH, inflammation, or necrosis separately. All measures of pathology showed no significant differences (*p*>0.05) in B6FVB TRAMP(+) liver tissues treated with either PBS (no therapy) control or four doses of 2.5 × 10^7^ CRC2631 every three days. P-values denote student t-test significance.

### CRC2631 preferentially colonizes primary tumors and metastases

Next, we sought to determine the *in vivo* tumor-targeting capability of CRC2631 in TRAMP animals.

First, we devised a strategy that not only permits longitudinal detection of CRC2631 in live-treated animals using fluorescence imaging, but also makes it possible to selectively isolate CRC2631 from harvested organs for quantitative bio-distribution assays.

The fluorescence reporter iRFP720 and a chloramphenicol resistance cassette were introduced into CRC2631, generating CRC2631^*iRFP720*-*cat*^. We replaced the kanamycin resistance cassette inserted into the Δ*thyA* deletion site with a gene fusion that constitutively expresses the iRFP720 fluorescent protein [32] and the *cat* chloramphenicol resistance cassette (Figure 3a). In comparison to CRC2631, CRC2631^*iRFP720*-*cat*^ produces visible iRFP720 fluorescence signal and grows in chloramphenicol media (Figure 3b, 3c), making it suitable for *in vivo* tumor-targeting studies.

**Figure 3.**
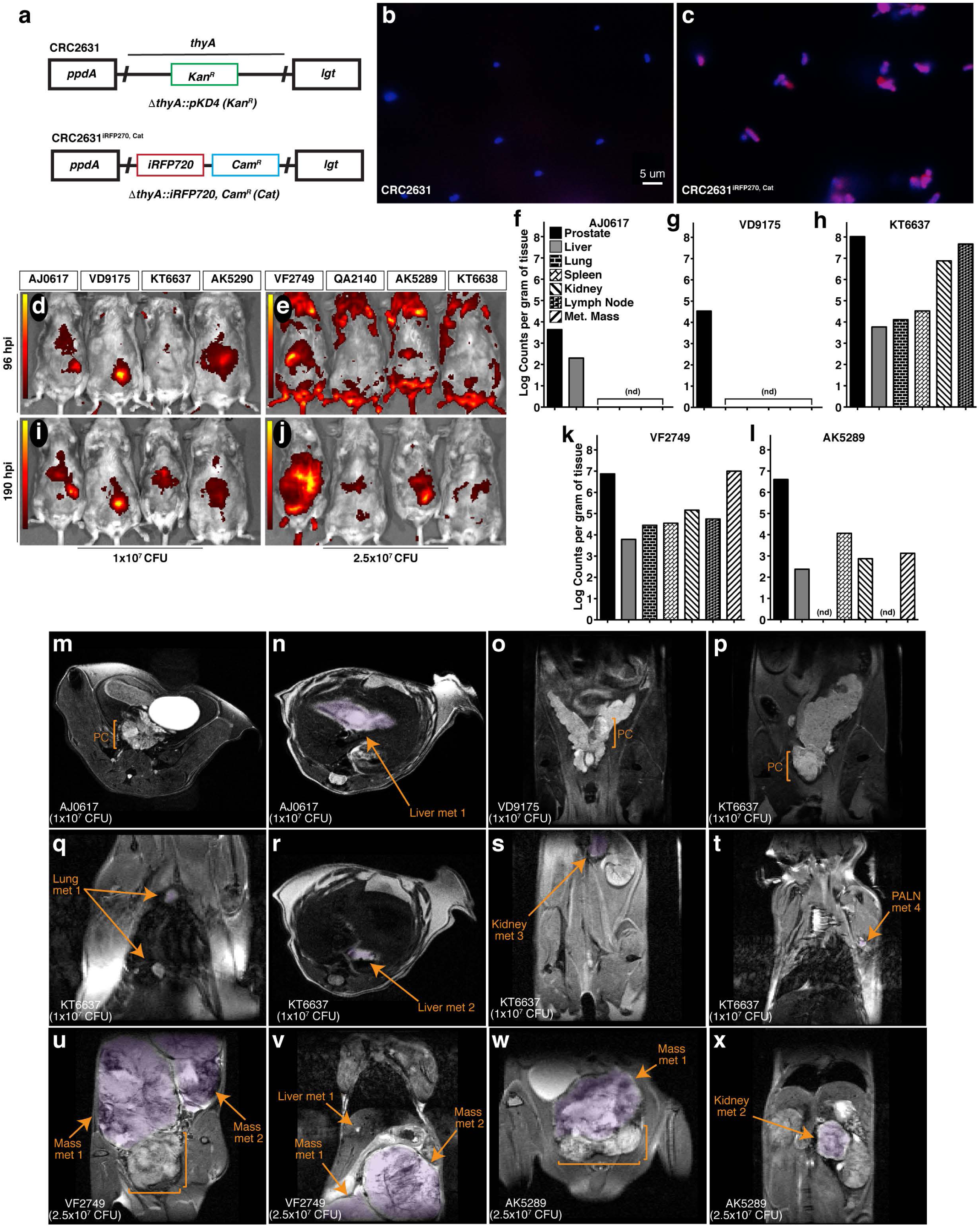
CRC2631 targets TRAMP primary and metastatic tumors. (a) Re-engineering strategy used to create CRC2631^*iRFP720*-*cat*^. The CRC2631 Δ*thyA*::Kan^R^ region (top) was replaced with genes that constitutively express iRFP720 and chloramphenicol (Cam^R^) (Cat) resistance (bottom) for fluorescent detection in TRAMP models *in vivo* and selective recovery of CRC2631^*iRFP720*-*cat*^ from tissue homogenates in biodistribution assays. (b, c) CRC2631^iRFP720-*cat*^ iRFP720 expression confirmed by microscopic examination of (b) CRC2631 and (c) CRC2631^iRFP720-*cat*^ mounted in Vectashield+DAPI stain and observed using a 63x objective with Cy5 and DAPI filter exposure overlay to detect iRFP720 and chromosomal DNA signal, respectively (red = Cy5 signal, blue = DAPI signal). (d-x) Analysis pipeline to assay CRC2631^*iRFP720*-*cat*^ tumor targeting tropism in the TRAMP model. Individual mice were assigned six-digit ID numbers [boxes above the in vivo fluorescent scan data (d-j)] to associate CRC2631^*iRFP720*-*cat*^ tumor targeting with unique metastatic burdens. (d-j) Two groups of >25 week-old B6FVB TRAMP(+) mice (*N=3*) were treated with (d, i) 1×10^7^ or (e, j) 2.5×10^7^ CRC2631^*iRFP720*-*cat*^. Using an IVIS *in vivo* fluorescent imaging system, fluorescent scans of live mice were conducted at (d, e) 96 and (i, j) 190 hours post injection. Images were spectrally unmixed against negative controls (AK5290, KT6638) to detect CRC2631^*iRFP720*-*cat*^-associated iRFP720 signal. Color bars indicate CRC2631^*iRFP720*-*cat*^ -associated iRFP720 signal intensity (red = low CRC2631^*iRFP720*-*cat*^-associated iRFP720 signal, yellow = high CRC2631^*iRFP720*-*cat*^-associated iRFP720 signal). (f-l) Enumeration of CRC2631^*iRFP720*-*cat*^ colony counts per gram tissue at 190 hours post injection with (f-h) 1×10^7^ and (k-l) 2.5×10^7^ CRC2631^*iRFP720*-*cat*^ as a direct measure of CRC2631^*iRFP720*-*cat*^ tissue targeting tropism. Enumerated tissue types are indicated in (f); Met Mass = discrete metastatic masses in the peritoneal cavity. (nd) = tissues with no detectable CRC2631^*iRFP720*-*cat*^/g tissue counts. (m-x) MRI scans of live B6FVB TRAMP(+) mice taken before tissue collection to confirm primary prostate tumor (PC, brackets) and metastatic tumor burden profiles of each mouse (metastatic purple highlighted regions indicated by arrows). Animal IDs and CRC2631^*iRFP720*-*cat*^ injection levels are indicated in the lower left of each MRI scan. (t) PALN = proper axillary lymph nodes. (u-v) Metastatic masses in animal VF2749 were attached to the right kidney in the upper peritoneal cavity. (w) Metastatic mass in animal AK5289 was adjacent to the primary prostate tumor. MRI-identified metastatic burdens in the TRAMP model correlate with CRC2631^*iRFP720*-*cat*^-associated tissue targeting signal identified by in vivo fluorescent scans of live mice (d-j) and CRC2631^*iRFP720*-*cat*^ colony enumeration from tissue samples (f-l) at 190 hours post injection.

Two groups (*N=*3) of B6FVB TRAMP(+) animals were scanned by MRI to establish tumor burden. All of the B6FVB TRAMP(+) animals exhibited prostate tumors and metastases to various visceral organs (Figure 3m-x); one animal had several metastatic masses in the peritoneal cavity (Figure 3u-v). To determine CRC2631^*iRFP720*-*cat*^ tumor-targeting capabilities, these animals received IV injections of 10^7^ or 2.5×10^7^ CFU of CRC2631^*iRFP720*-*cat*^. One additional B6FVB mouse was included in each dose group (AK5290 and KT6638 for the 10^7^ and 2.5×10^7^ CFU groups, respectively) as a fluorescence background control. We first determined CRC2631^*iRFP720*-*cat*^ bio-distribution at 96 hours and 190 hours post injection (hpi) by detecting the iRFP signal of CRC2631^*iRFP720*-*cat*^ at the indicated time-points using the fluorescence *in vivo* imaging system (IVIS). In both dosage groups, we detected above background iRFP signals in two of the three animals treated with CRC2631^*iRFP720*-*cat*^. We detected high intensity iRFP foci in the prostate and peritoneal cavity regions at 96hpi (Figure 3d, 3e). These signals coincided with the positions of primary and metastatic legions in from MRI images and persisted for over three days (190hpi) (Figure 3i, 3j), suggesting that CRC2631^*iRFP720*-*cat*^ preferentially colonizes tumor tissues. To test this hypothesis, we determined CRC2631^*iRFP720*-*cat*^ load in tumor tissues (prostate and bulk metastases), blood, lung, lymph nodes, liver, spleen, and kidneys harvested from the 190hpi animals above (detection thresholds for the aforementioned tissues were 6.58×10^1^, 1×10^3^, 6.73×10^3^, 4.86×10^4^, 2.63×10^2^, 9.69×10^3^, and 4.70×10^2^ CRC2631^*iRFP720*-*cat*^ counts per gram tissue, respectively). The indicated tissues were harvested and their respective counts of CRC2631^*iRFP720*-*cat*^ per gram of tissue were determined under chloramphenicol selection (see methods). Detectable CRC2631^iRFP720-*cat*^ was enriched in the prostate at the 10^7^ and 2.5×10^7^ CFU dosing levels (Figure 3f-l). Two out of three animals in the 10^7^ CFU group had detectable colonies only from the prostate and liver tissues. One animal in the 2.5×10^7^ CFU group (i.e., QA2140) did not yield any colony in the analyzed tissues; however, the remaining two animals (VF2749 and AK5289) showed higher colony counts in the prostate and bulk metastases compared to the remaining tissues (Figure 3k-l). No detectable CRC2631^iRFP720-*cat*^ counts were present in whole blood. Comparing bacterial load in the liver versus in tumor tissues provides a measure of tumor-targeted bacterial colonization. The prostate to liver ratio of CRC2631^*iRFP720*-*cat*^ counts ranged from 20:1 to 18000:1 and 1220:1 to 1690:1 in the 10^7^ CFU and 2.5×10^7^ CFU treated animals, respectively (Figure 3f-l). Two (VF2749 and AK5289) of the three animals in the 2.5×10^7^ CFU dose group exhibited several prominent metastases (Figure 3u-w) and the metastases to liver CRC2631^*iRFP720*-*cat*^ count ratio ranged from 1640:1 to 2990:1 (Figure 3k-l). Taken together, these data indicate that CRC2631^*iRFP720*-*cat*^ targets primary tumors and metastases.

### CRC2631 is genetically stable inside the host

The genetic alterations that attenuate CRC2631 and contribute to its tumor-targeting capability are permanently integrated in its genome. This reduces the likelihood that CRC2631 will regain toxicity and/or lose its tumor targeting capability due to *de novo* mutations inside the host environment; however, it remains a possibility. To determine the genetic stability of CRC2631 inside the host, we performed longitudinal whole genome sequencing and short nucleotide polymorphism (SNP) analyses of CRC2631 prior to treatment and tumor-passaged CRC2631 in B6 TRAMP(+) mice. Animals (*N=4*) were treated intravenously with CRC2631 (2.5×10^7^ CFU), and then CRC2631 was isolated from prostate tissues harvested at 96 or 190 hpi. We recovered three isolates from the 96 hpi (CRC2631a-c) and one isolate from the 190 hpi (CRC2631d) prostate tissues (Figure 4a). We isolated DNA and performed Illumina Next Generation Sequencing (NGS) on 0 hpi CRC2631 (prior to treatment) and all tumor-passaged isolates to identify SNP mutations occurring in the host environment. Comparisons of the 96hpi or 190hpi versus the 0hpi sequences identified a total of two and three SNPs at 96hpi and 190hpi, respectively. The three 190hpi SNPs include the same two SNPs identified at 96hpi. To map these SNPs to specific genes we annotated genome information from the *Salmonella enterica* serovar Typhimurium LT2 strain (GenBank: AE006468.2) and its associated pSLT plasmid (GenBank: AE006471.2) [9]. There is no annotated genome information currently available for CRC2631 and CRC2631 is a direct derivative of LT2. With the exception of one SNP that mapped to an intergenic region, all of the remaining variants represent synonymous SNPs. The two 96hpi SNPs mapped to two distinct positions in the STM1021 locus, which is similar to the Gifsy-2 lysogenic bacteriophage region (*ninG*) in the LT2 genome (Figure 4b-c). The unique 190 hpi SNP consisted of a six base-pairs deletion (CCTGTT) in an intragenic region between pSLT064 and *ssbB* of the LT2-associated plasmid (pSLT) (Figure 4d).

**Figure 4.**
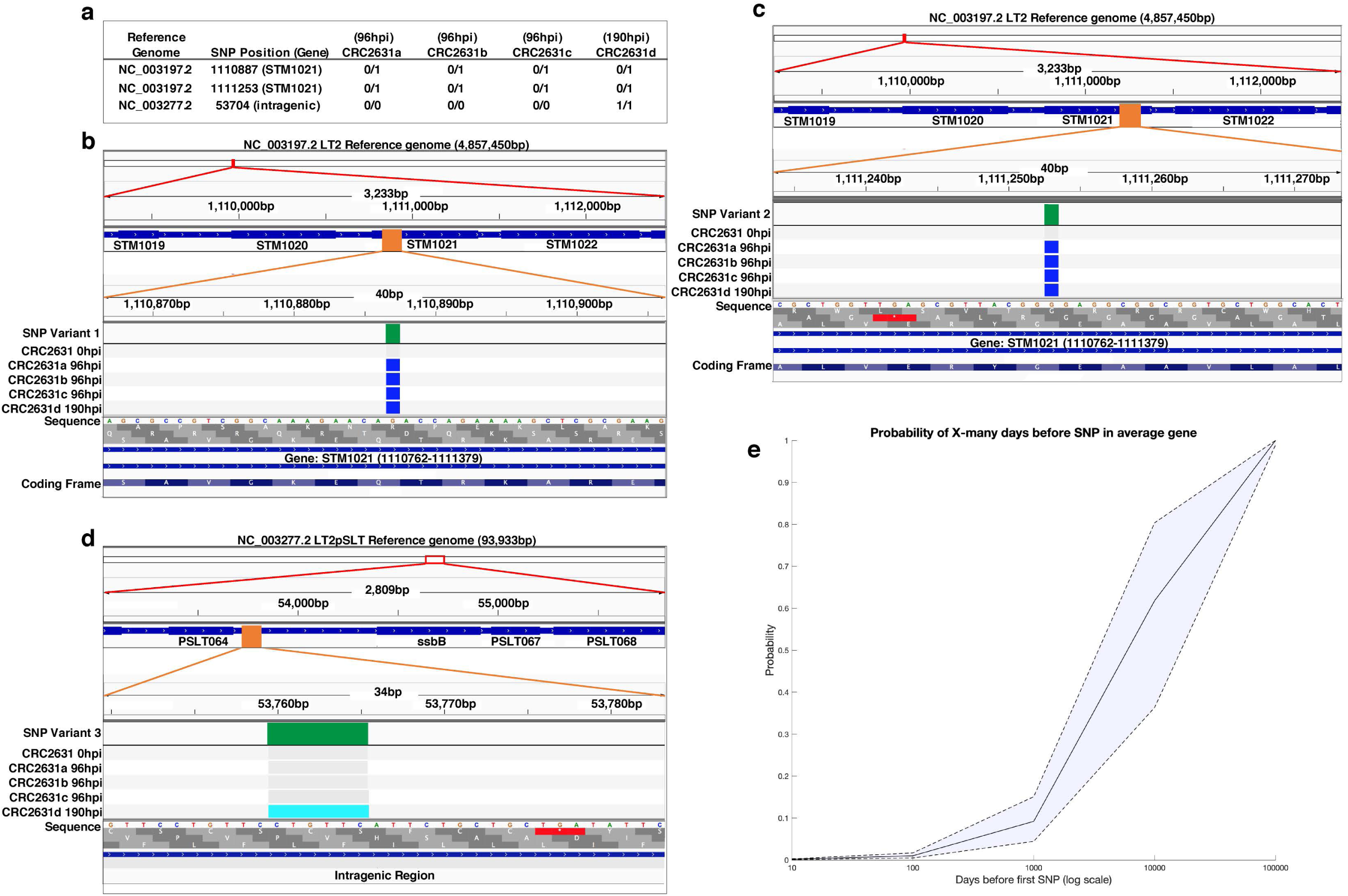
CRC2631 is genetically stable in tumors. (a) Short nucleotide polymorphisms (SNPs) identified in the tumor-passaged strains (CRC2631a-d) compared to the CRC2631 injection strain. Reads were assembled and mapped against the CRC2631 parental LT2 and associated stable pSLT plasmid sequences. SNPs unique to tumor passage were identified by sequencing CRC2631 injection aliquots (“CRC2631 0hpi”) before injection and CRC2631 isolated from individual B6 TRAMP(+) mouse prostate tumor tissue samples at 96 (CRC2631a-c) or 190 (CRC2631d) hours post injection. “0/0” indicates no SNP mutation in daughter strain compared to CRC2631 injection strain; “0/1” indicates a SNP mutation in daughter strain compared to CRC2631 injection strain; “1/1” indicates a SNP mutation in CRC2631 daughter strain that reverts back to the original LT2 sequence compared to CRC2631 injection strain. (b-d) Graphical representations of SNP locations accumulated in the B6 TRAMP(+) mouse prostate tumor environment over 96 and 190 hours using Integrative Genomics Viewer (v2.8.0). SNP locations are shown at three genomic resolutions. From top to bottom, the SNP location is indicated as a red box at the cytologic overview, followed by increase in the genomic resolution to the local gene region showing labeled gene coding regions as blue boxes and SNP location as an orange box, and finally showing the SNP location as a green box at nucleotide resolution. Green: Location of SNP mapped to the GenBank reference sequence. Grey: No change from CRC2631 parent. Blue: SNP mutation from CRC2631 parent. Light blue: deletion of 6 bp repeat in CRC2631 that reverts CRC2631d to original LT2 sequence. (e) SNP prediction modeling displaying the probability of an average gene in CRC2631 accumulating a first SNP after a given number of days in the tumor environment on a logarithmic scale. The probability of an average gene accumulating a first SNP after 10, 100, 1000, 10000 and 100000 days is 0.0015, 0.01, 0.0921, 0.6181, and 0.9999 respectively.

The observed low SNP frequency in tissue-passaged CRC2631 over an ∼8 day period argues that CRC2631 is genetically stable within the host. To evaluate the robustness of CRC2631 genetic stability, we first estimated the time it would take for CRC2631 to experience a SNP in any gene of interest. The LT2 lineage of the CRC2631 genome (including the stably associated pSLT plasmid) consists of ∼4951383 base pairs organized in ∼4548 predicted gene coding sequences. The average size of the gene coding sequences is 943.89bp [9]. Considering the observed SNP frequency rate (51.83±7.67 hours/SNP), it would take ∼9375 days for CRC2631 to acquire a SNP in any specific gene by chance. In a complementary approach, we modeled the risk probabilities of such an event over time (see methods). Our model predicts that the probability of an average gene accumulating a first SNP after 10, 100, 1000, 10000 and 100000 days to be: 0.0015, 0.01, 0.0921, 0.6181, and 0.9999, respectively (Figure 4e). The probability that CRC2631 will gain 0, 1, 2, 3, 4, and 5 mutations four days after treatment is predicted to be 0.21, 0.29, 0.23, 0.14, 0.07, and 0.03, respectively. Thus, CRC2631 is a stable tumor-targeting biologic.

### CRC2631 and checkpoint blockade combination reduces metastatic incidence

The observations that CRC2631 stably colonizes tumors, including metastases, prompted us to explore the possibility that CRC2631 reduces tumor burden in TRAMP animals. We previously reported that low doses of CRC2631 (10^7^ CFU administered weekly) modestly reduced prostate tumor size in TRAMP animals [8]. We first asked whether CRC2631 generates a more robust therapeutic response at a higher CRC2631 dose. Pre- and post-treatment MRI images were used to compare tumor size in response to therapy. Groups (*N=12*) of 8-10-week-old B6FVB TRAMP(+) animals were treated with PBS (control) or 2.5 ×10^7^ CFU CRC2631 administered IV every three days for a total of four treatments. We scored prostate tumor size in control versus CRC2631-treated animals 21 days after treatment initiation and found that CRC2631 did not significantly reduce prostate tumor size, compared to the PBS control (Figure 5a; *p*<0.6799, 30.77mm^3^±76.07 versus 42.30mm^3^±52.79 respectively). CRC2631 targets and directly kills murine and human prostate cancer cells *in vitro* (Supplemental Figure 2), raising the possibility that unknown resistance mechanisms protect tumor cells from CRC2631-mediated cell death *in vivo*. These inhibitory signals may be tumor cell-intrinsic and/or involve the tumor immune microenvironment.

**Figure 5.**
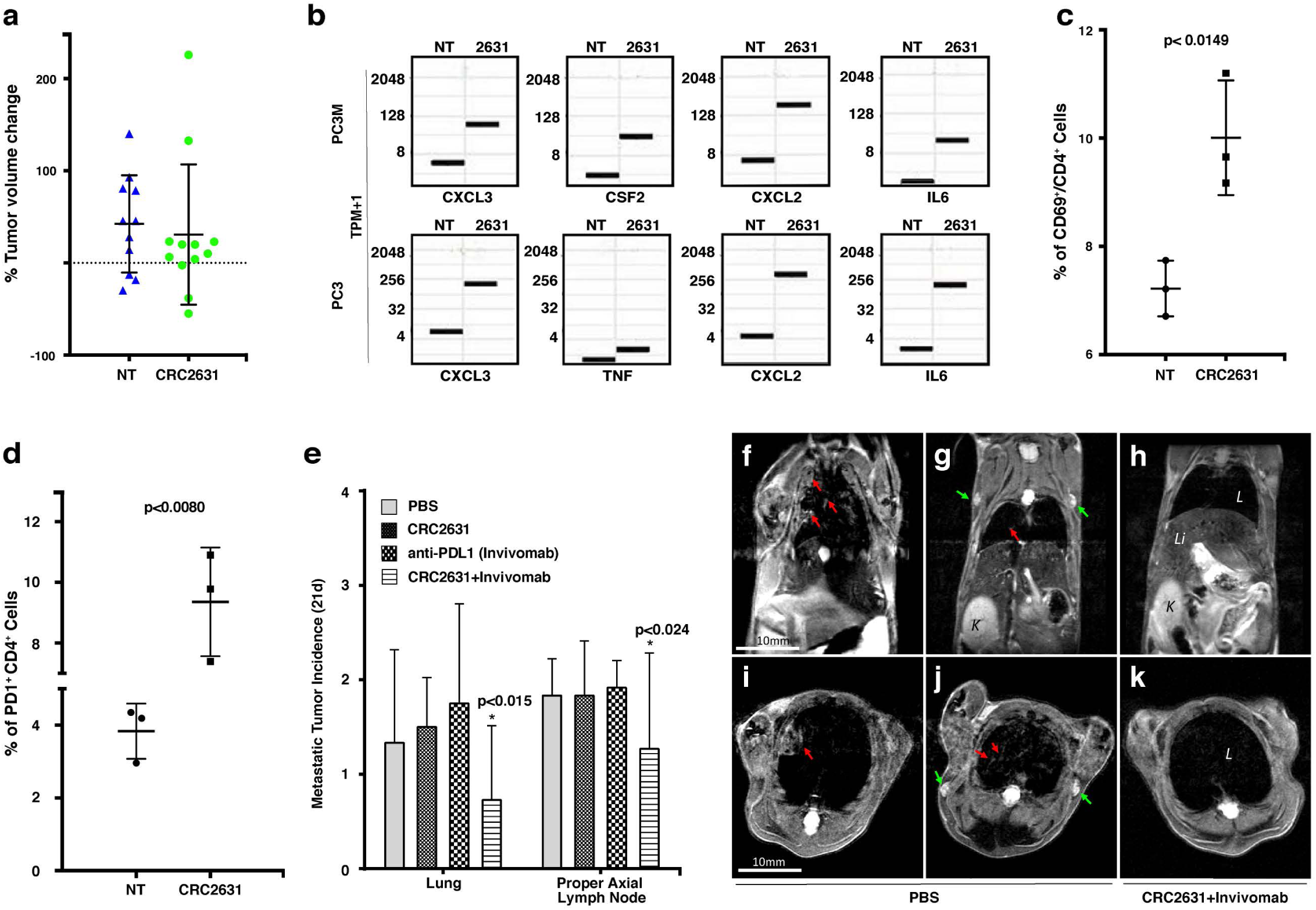
CRC2631/PD1 blockade combination treatment reduces metastatic burden. (a) Graph showing percentage change of ventral prostate tumor volume in 8-10-week-old B6FVB TRAMP(+) animals (*N=12*) following IV treatment with PBS [no treatment (NT)] or 2.5 × 10^7^ CRC2631. Animals received a single dose of indicated treatment every three days for a total of 4 injections. Tumor volume was determined using the volumetric image analysis platform IMARIS BITPLANE and MRI images taken 5-7 days before and 21 days after treatment. (b) Box plot showing gene expression change in Transcript Per Million on a log2 scale (TPM+1, *N=3*) in PC3M and PC3 human prostate cancer cell lines before and after treatment with CRC2631. NT = No Treatment, 2631 = CRC2631 treated cells. (c, d) Flow cytometric profiling of tumor-infiltrating lymphocytes (TIL) in metastasized lymph nodes extracted from PBS [no treatment (NT)] or CRC2631-treated B6FVB TRAMP(+) animals. Cells were sorted on CD3, CD69, CD4 (c), and CD3, CD4, PD1 (d). Graph depicts the frequency of the indicated TIL phenotype across groups. P-values are derived from students’ t-test analyses. (e) Enumeration of metastases observed in lymph nodes and lung tissues from MRI images taken 21 days after treatment from groups of 8-10 weeks old B6FVB TRAMP(+) animals (*N=12*) treated (IV) with PBS or 2.5 × 10^7^ CFU of CRC2631 or 0.5 mg of a murine anti-mouse PDL1 antibody (Invivomab), or a combination of CRC2631 (2.5×10^7^ CFU) and Invivomab (0.5 mg). Animals in each group received a single dose of the indicated treatment every three days for a total of four injections. P-values denote student t-test significance. (f-k) Representative *in vivo* MRI images of B6FVB TRAMP(+) mice 21 days after start of CRC2631+Invivomab treatment (h, k) compared to PBS control group (f-g, i-j). Red arrows: lung metastasis; green arrows: lymph node metastasis. L: lungs; Li: liver; K: kidney.

We turned our focus to an interaction between CRC2631 and immune cells and asked whether tumor-targeted CRC2631 generates an anti-tumor immune response that tumors rapidly inhibit via immune checkpoint mechanisms. Tumor cells express program death ligand-1 (PDL1), which interacts with PD1 on the surface of immune T-cells to inhibit anti-tumor immune activities [33, 34].

RNA sequencing data shows an upregulation of immunogenic chemokines and cytokines (CXCL, CSF2, IL6, and TNF) in human prostate cancer cells (PC3, PC3M) (Figure 5b) and Luminex-assayed cytokine response in mice treated with CRC2631 (Figure 1i-j). Similar to men, TRAMP animals develop a robust immunosuppressive microenvironment in prostate tumors [30, 35-37], which will likely mask the effect of CRC2631. Thus, we focused on distant metastases to determine whether metastases-targeted CRC2631 recruits and activates tumor-infiltrating lymphocytes (TILs) *in vivo*. TRAMP animals develop unambiguous lymph node and visceral organ metastases that can be readily detected in MRI images [[23], Figures 2m-x]. Lymph nodes containing MRI-verified metastases were harvested either from CRC2631-treated or PBS control TRAMP animals. Flow cytometric analyses showed that CRC2631 significantly elevates the frequency of activated CD4^+^ (CD69^+^/CD4^+^) TILs in metastasized lymph node tissues (Figure 5c). Consistent with this, others have shown that *aroA* deletion augments bacterial immunogenicity [38]. CD4^+^ TILs mediate an anti-cancer immune response by activating tumoricidal CD8^+^ T-cells [39]. In contrast, regulatory T-cells (T^reg^) suppress CD8^+^ T-cells [40, 41]. CRC2631 treatment did not enhance T^reg^ frequency compared to PBS control tissues (data not shown). Interestingly, histological analyses did not show an increase of CD8^+^ TILs in metastasized lymph nodes or lungs derived from the therapy animals compared to controls (data not shown), suggesting T-cell exhaustion. Congruently, CRC2631 increased the proportion of CD4^+^ TILs expressing the exhaustion marker PD1 (CD4^+^ PD1^+^) (Figure 5d). Targeted inhibition of the PDL1-PD1 signaling axis restores anti-cancer immune activity and generates significant clinical benefits in patients with immunogenic cancers [41, 42]. Our observations suggest that CRC2631 may reduce tumor burden by enabling an anti-tumor immune response in the PDL1/PD1 blockade setting.

Eight-to-ten-week old animals (*N=12*) were scanned by MRI to control for tumor burden across groups. These animals were treated with 200uL of PBS (vehicle control) or CRC2631 (2.5×10^7^ CFU) or murine anti-PDL1 antibodies (Invivomab, 0.5mg) or a cocktail of CRC2631-Invivomab. The dosing regimen consisted of a single dose of the indicated treatment every three days for a total of four infusions. Post-treatment lung and lymph node MRI images were used to enumerate and compare metastasis incidence across groups at 21 days after treatment (Figure 5e-f). PBS control animals showed an average of 1.83±0.389 and 1.33±0.985 metastases in proper axial lymph nodes and lungs, respectively. Alone, CRC2631 or Invivomab treatments did not significantly reduce metastasis to the lymph nodes or the lung. In contrast, the CRC2631-Invivomab combination treatment reduced metastasis incidences in lymph nodes and the lung. CRC2631-Invivomab averaged 1.27±1.01 proper axial lymph node metastatic incidences after 21 days compared to 1.83±0.577 in CRC2631 and 1.92±0.289 in Invivomab alone. In the lung, CRC2631-Invivomab showed an average of 0.727±0.786 metastases compared to 1.50±0.522 in CRC2631 and 1.75±1.06 in Invivomab alone. Thus, CRC2631-checkpoint blockade combination treatment reduces metastatic burden.

## Discussion

Conventional chemotherapies are not cancer-specific and as a result generate significant morbidities. Several toxicity-mitigating strategies have been proposed, including the use of genetically attenuated bacteria that specifically colonize tumor tissues to deliver therapeutics [43]. However, the lack of bacterial cancer targeting (BCT) strains that are objectively safe continues to limit the clinical utility of these technologies. This is partly because preclinical tumor-targeting and safety evaluations have relied on moderate cancer models in immune suppressed animals [4-6]. The most studied BCT strain, VNP20009, safely colonized tumors in immune-suppressed animal models but fails to generate a therapeutic signal in human patients, presumably because of rapid immune clearance by the host [7]. Here, we describe the toxicological, tumor-targeting, and therapeutic profiles of CRC2631 in a syngeneic mouse model of aggressive prostate cancer (TRAMP). We show that CRC2631 is a safe and genetically stable biologic that persistently colonizes tumors, including metastases.

Comparing the toxicity and tumor-targeting profiles of CRC2631 against those of VNP20009 showed that VNP20009 generates more toxicity than CRC2631 and poorly targets tumor tissues in immune-competent TRAMP animals (Supplemental Figure 3b-3d). Consistent with these observations and in contrast to earlier findings from nude animals, a single injection of 2×10^4^ CFU VNP20009 also showed significant liver toxicity and poor tumor targeting capabilities in an immune-competent mouse model of mammary carcinoma [31], whereas CRC2631 exhibited no significant liver toxicity after three orders of magnitude higher injections of CRC2631 into the immune-competent B6FVB TRAMP model (Figure 2).

CRC2631 partly owes its tolerability and enhanced tumor-targeting characteristics to its unique genomic evolution. CRC2631 was isolated from a collection of naturally occurring mutant strains that arose after maintaining the Salmonella LT2 in nutrient-limiting conditions for over four decades. This long-term selection generated a diverse array of genetic alterations while removing the selective pressure to maintain factors that are required for bacterial virulence in a human host. In addition, CRC2631 is deficient in lipid polysaccharide biosynthesis, leading to even less toxicity. Furthermore, CRC2631 is auxotrophic for aromatic amino acids and thymine, favoring CRC2631 growth specifically in metabolically rich environments such as cancers. These properties not only augment its tumor targeting but also limit its toxicity. Consistently, CRC2631 is specifically enriched in tumor tissues (Figure 3), and does not cause overt toxicity (Figure 2). Additional support for CRC2631 safety and preferential colonization of tumor tissues comes from our findings that CRC2631 is well-tolerated in healthy dogs. Serial blood analyses revealed relatively normal organ function (Supplemental Figure 4).

In addition to the preferential growth in cancers, other mechanisms likely contribute to CRC2631 tumor tropism. Kasinskas and Forbes (2006) show that Salmonella strain SL1344 requires wild-type serine, aspartate, and ribose chemoreceptors for active targeting of colon carcinoma cylindroids *in vitro* [44]. Additionally, we screened CRC2631 against a library of human cell surface glycoproteins to identify specific cell surface molecules required for CRC2631-host interaction. We found that CRC2631 binds to mannose-linked terminal disaccharides surface glycoproteins 10-to >400-fold more efficiently than to glycoproteins lacking mannose-linked terminal disaccharides (unpublished data). Glycoproteins that CRC2631 bound with high affinity are commonly found on cancer cells [45]. This suggests that cancer-specific surface molecules promote the selective entry of CRC2631 into cancer cells.

Longitudinal genome analyses of tumor-passaged CRC2631 showed that CRC2631 remains genetically stable within the tumor microenvironment. At the 2.5×10^7^ CFU dose, mutation rate modeling estimates a 0.15% probability that an average gene within CRC2631 will acquire a mutation inside the host within ten days of treatment. An average gene within CRC2631 will require 100,000 days inside the host to reach the absolute certainty that it will acquire a SNP, which is well beyond the time window of any therapy. We note that a limitation of our modeling approach is that it makes predictions for an average gene within CRC2631 and does not take into account base pair level information for individual genes. Future work could extend the model to this level but doing so would also require larger samples over deep time points. These modeling data allow one to rationally assign risk levels for specific dosing regimens in other pre-clinical models or in human patients. Collectively, our findings indicate that CRC2631 is a genetically stable biologic that safely targets tumors, including metastases, in immune-competent hosts.

Tumor-localized CRC2631 fails to reduce the size of primary prostate tumors in TRAMP animals. This could be due to a sub-optimal intra-tumoral CRC2631 load; a higher and safe dosing regimen and/or direct intratumoral CRC2631 delivery may be considered. This result also could be due to the aggressiveness of the TRAMP model. In contrast to other mouse cancer models where oncogenesis is triggered in a limited number of cells over a defined time interval, androgen-driven expression of SV40 antigens transforms prostatic tissues more broadly and continuously, leading to sustained and rapid tumor overgrowth. This may potentially mask a CRC2631 tumor suppressive effect. Prostate cancers progress more slowly in human patients. Expanding the evaluation of CRC2631 therapeutic profile to other tumor models will be informative.

Importantly, CRC2631 reduced metastasis incidence in the setting of checkpoint blockade. This is significant because metastasis is the main cause of cancer-associated deaths and no effective immunotherapy against prostate cancer currently exist.

## Materials and Methods

### Growth of bacterial cultures

See Supplemental Table 1 for list of bacteria used in this study. Isolated colonies of bacteria were grown from −80 ^°^C stock aliquots frozen in 25% glycerol (Fisher) on solid or liquid LB media (Fisher) containing 200 µg/ml thymine (Arcos Pharmaceuticals) and selective antibiotics [50 µg/mL kanamycin (Sigma), 50 µg/mL ampicillin (Sigma), or 20 µg/mL chloramphenicol (Gold Biotechnology)] as required. Bacteria grown on solid media was incubated for 24-30 h at 37 °C before use. Liquid media cultures were incubated in 50 mL sterile tubes for 20-24 h in a 37 °C, 220 rpm dry shaking incubator. Strains grown for injection were washed with sterile phosphate buffered saline (PBS) (Rocky Mountain Biologicals) and concentration adjusted for injection (Supplemental Figure 1) and for *in vitro* cell viability assays.

### Cell lines and cell culture conditions

See Supplemental Table 1 for list of cell lines used in this study. All cell lines were obtained from ATCC (Manassas, VA). The RWPE-1 cell line was maintained in Keratinocyte Serum Free Medium (K-SFM) media (Gibco); PC3 cells were grown in Ham’s F-12K Medium (Gibco) supplemented with Fetal Bovine Serum to a final concentration of 10%; and PC3M cells were maintained in RPMI 1640 (Gibco) supplemented with 10% FBS, 1mM sodium pyruvate (Fisher), 1X non–essential amino acids (Thermofisher) and 2mM L-Glutamine (Fisher). TRAMP-C2 cells were grown according to ATCC guidelines. All cells were maintained at 37 ^°^C with 5% CO_2_.

### Construction of CRC2631^*iRFP720*-*cat*^

The standard Datsenko and Wanner recombination protocol [46] was used to engineer the Δ*thyA*::P^tac^-*iRFP720 cat* (Cam^R^) insert into CRC2631, replacing the Kan^R^ gene cassette at the Δ*thyA*::pKD4(Kan^R^) deletion site to create CRC2631^*iRFP720*-*cat*^ (Figure 3a). A 50 bp of flanking homology upstream and downstream of the region internal to the CRC2631 Δ*thyA*::pKD4 (Kan^R^) insertion was engineered using a megaprimer primer construct to replace the Kan^R^ gene cassette with a wild-type P^tac^ promoter, the *iRFP720* gene from pBAD/HisB-*iRFP720* (Addgene) [32], and the wild-type *cat* (Cam^R^) gene from pRE112 [47]. An overnight liquid culture of CRC2631pKD46 was grown in 10mL of LB+200 µg/ml thymine, 50 µg/mL ampicillin and 50 µg/mL kanamycin at 30°C, 220 rpm dry shaking incubator. A 0.25 mL overnight CRC2631pKD46 culture (1% inoculum) was sub-cultured into 25mL LB+200 µg/ml thymine, 50 µg/mL ampicillin, 50 µg/mL kanamycin and 100mM L-arabinose (Sigma) media and grown in sterile 50 mL tubes on a 30°C, 220rpm dry shaking incubator. After 10 h, cells were recovered by 10 min centrifugation at 4000 rpm and washed 4 times in 1 mL cold sterile water, then resuspended in 75 µl sterile 10% glycerol. Using a 0.2 cm electroporation gap cuvette (Fisher) 1 µg of insert DNA was electroporated (2.5 kV) (Electroporator 2510, Eppendorf) into the 10% glycerol CRC2631pKD46 cell suspension. One mL of LB+200 µg/ml thymine was added to the cuvette and the cells were allowed to recover at 37°C for three hours. The cells were centrifuged at 13.2 k rpm for 1 min, the supernatant discarded, and resuspended in 500 µl LB +200 µg/ml thymine, then spread on selective plates containing LB +200 µg/ml thymine and 7.5 µg/mL chloramphenicol. These plates were incubated for 24 h at 37 °C to recover Cam^R^ Kan^S^ transformants. The temperature-sensitive pKD46 helper plasmid was lost by overnight growth at 42 °C, growth of 20-200 isolated colonies on LB +200 µg/ml thymine +20 µg/mL chloramphenicol plates, and the target Cam^R^, Kan^S^, Amp^S^ antibiotic resistance profile confirmed using replica plating. The resulting Cam^R^, Kan^S^, Amp^S^ Δ*thyA*::P^tac^-*iRFP720 cat* (Cam^R^) insertion CRC2631^*iRFP720*-*cat*^ construct was confirmed by PCR analysis and fluorescence microscopy (Figure 3b-c).

### Fluorescence microscopy of CRC2631^*iRFP720*-*cat*^

CRC2631^*iRFP720*-*cat*^ was grown for 24 h at 37 °C, 220 rpm in LB +200 µg/mL thymine +20 µg/mL chloramphenicol, washed in one volume of PBS, fixed in one volume of PBS+4% paraformaldehyde, washed in one volume PBS, mounted under a glass coverslip at a 1:1 ratio in Vectashield+DAPI (Vector Laboratories) stain, cured in the dark at room temperature for 2 h, sealed, then observed on a Zeiss Axiovert 200M fluorescent microscope using a 63x objective with 1.4NA. A Hamamatsu Orca-ER monochrome CCD camera was used to take 900 ms Cy5 filter + 24 ms DAPI filter exposures, which were pseudocolored and overlaid to confirm fluorescent detection (Figure 3c).

### Mice

See Supplemental Table 1 for mouse genotypes used in this study. **Tr**ansgenic **A**denocarcinoma of **M**ouse **P**rostate (TRAMP) mice of purebred C57BL/6-Tg(TRAMP)8247Ng/J (B6) (Jax Laboratories) or hybrid C57BL/6-Tg(TRAMP)8247Ng/J x FvBNHsd (Envigo) (B6FVB) background were genotyped at 21-28 days of age to distinguish between heterozygous TRAMP(+) animals positive for the PB-Tag SV40 oncogene and TRAMP(-) animals negative for the PB-Tag SV40 oncogene as previously described [48]. B6 and B6FVB TRAMP mice were allowed to grow to 8-31 weeks of age before use in studies. Food [LabDiet5001 (LabDiet) or AIN-93M (Research Diets)] and water were provided *ab libitum*. Animals were observed and weighed on a daily basis during all studies. All animal studies were conducted in accordance with the principles and procedures outlined in the National Institutes of Health Guide for the Care and Use of Animals under the University of Missouri Animal Care and Use Committee supervision (MU IACUC protocols #8602 and #9501).

### MRI imaging

All mice used in toxicity studies were imaged on a Bruker AVANCE III MRI platform. This system has the capability of achieving a 50 µm resolution for imaging tumor models. Mice were anesthetized using 3% isoflurane; anesthesia was maintained with 0.5-2% isoflurane to keep breathing rate at 30 bpm during which axial and coronal scans of the mouse body were performed. Images were taken using ParaVision 6 software (Bruker BioSpin Inc). Prostate tumor volumes were measured using Imaris software (Bitplane) to normalize injection groups for an average range of primary tumor burden and to measure therapeutic response to treatment.

### Toxicological studies

All B6FVB TRAMP(+) mice used in toxicity studies were scanned using a Bruker AVANCE III MRI platform as described above to confirm tumor burden. Mouse tumor burdens were graded by size using the Imaris software package. Mice were randomly assigned to study groups ensuring that each group had a representative range of tumor burden. Mice groups were injected interperitoneally with up to 5 × 10^7^ CRC2631 or VNP20009 or sterile PBS carrier volume (100-500 µl) or intravenously (tail vein) with 2.5 × 10^7^ CRC2631 in 200 µl PBS four to fourteen times with a weekly or three-day interval between doses until study completion or until loss of 50% of the group, after which tumor burden was determined using MRI scans and the mice subsequently evaluated for life extension.

### Cytokine assays

Whole blood samples were taken from B6FVB TRAMP(+) mice via saphenous vein draw into capillary tubes containing EDTA anticoagulant (Ram Scientific) 2 h before and 2 h after first CRC2631 or VNP20009 injections to measure the innate CRC2631 and VNP20009 inflammatory cytokine response. Blood was placed on ice and plasma immediately extracted from the whole blood by centrifugation for 10 min at 1000 x g in a 4 °C centrifuge followed by transfer of the supernatant to a new Eppendorf tube. Platelets were removed from the supernatant by centrifugation for 15 min at 2000 x g in a 4 °C centrifuge. The resulting plasma supernatant was transferred to a sterile Eppendorf tube and stored at −80 °C until the cytokines were measured using a Milliplex xMAP Mouse High Sensitivity TCell Magnetic Bead Panel kit MHSTCMAG-70K (Millipore) following the kit protocol on a Luminex 200 system with Xponent (v2.7). Data analysis was performed using Analyst (Millipore).

### *In vivo* fluorescent imaging

All mice were fed a defined AIN-93M Mature Rodent diet (Research Diets) for a minimum of seven days to minimize feed-related autofluorescence in the gastrointestinal system [49]. B6 or B6FVB TRAMP(+) animals were injected either intraperitoneally with 1 × 10^6^ CRC2631 pRSTmCherry or VNP20009 pRSTmCherry, or injected intravenously (tail vein injection) with 1 × 10^7^ or 2.5 × 10^7^ CRC2631^*iRFP720*-*cat*^. Fluorescent imaging was performed using a Xenogen IVIS 200 *in vivo* fluorescence system (Perkin-Elmer) and analyzed using Living Image software (v4.7.3). iRFP720 expression spectral unmixing was performed as previously described to detect CRC2631^*iRFP720*-*cat*^ associated iRFP720 *in vivo* [32]. Images containing mCherry RFP (Supplemental Figure 3) were spectrally unmixed to distinguish the CRC2631 or VNP20009-associated signal *in vivo* using mCherry spectral unmixing settings in the Living Image software.

### Biodistribution analysis

B6FVB TRAMP(+) mice were injected intravenously (tail vein injections) with 200 µl sterile PBS containing 1.0 × 10^7^ or 2.5 × 10^7^ CRC2631^*iRFP720*-*cat*^. Mice were euthanized 190 hours post injection. Whole blood, lung, liver, spleen, kidneys, prostate, and proper axial lymph nodes as well as any discrete metastatic tumor masses were collected, weighed, and kept on ice. Whole blood samples were immediately diluted 1/10 in 25% glycerol and PBS and stored at −80 °C. Tissue samples were homogenized in 3 mL sterile PBS for 20 seconds on ice using a TissueRuptor homogenizer (Qiagen) with sterile tips, mixed with 3 mL of sterile 50% glycerol, and stored at −80 °C. All tissue samples were later thawed, passed through 40 µm sterile filters (BD Biosciences) and immediately diluted, spotted in triplicate on selective LB +200 µg/ml thymine +20 µg/mL chloramphenicol plates, incubated at 37 °C and enumerated after 24 h following the Miles and Misra method [50].

### Histopathological analyses

Histopathological analyses were performed on samples obtained at necropsy from 33-week-old male B6FVB TRAMP(+) mice (four treated with CRC2631, four untreated) to examine the effects of CRC2631 administration on liver tissues. The four 31-week-old treated mice were given four intraperitoneal injections of 2.5 × 10^7^ CRC2631 at three-day intervals, and the untreated mice were given 250 µl sterile PBS intraperitoneal injections. Animals were euthanized at the study endpoint (end of week 33) by CO_2_ asphyxiation and subsequent exsanguination. The liver tissues were collected and immediately fixed in 10% buffered formalin (Fisherbrand), paraffin embedded, sectioned (5 µm thick sections), mounted on glass slides and stained with hematoxylin and eosin for histopathologic examinations by light microscopy. The liver tissue pathology grading system evaluating necrosis, inflammation, and EMH as measures of pathology was performed as previously described [31].

### Flow cytometric analysis of infiltrating lymphocytes

Metastasized lymph nodes were homogenized, and cells were isolated. Immune cell phenotypes were determined via flow cytometry with a FACSAria (BD Biosciences) using following antibodies: anti-mouse CD3 FITC, anti-mouse CD4 PE, anti-mouse CD8 PerCP/Cyanine 5.5, anti-mouse CD69 APC, anti-mouse PD1 BV421. All antibodies were purchased from Biolegend.

### RNA isolation and RNAseq

Prostate cells, benign (RWPE-1) and cancer (PC3 and PC3M) at 80% confluency, were treated with 10^4^ CFU of CRC2631 for 1.5 h at 37 ^°^C and 5% CO_2_. Total RNA was isolated from CRC2631-treated and non-treated samples using the RNeasy Plus kit (Qiagen) as per the manufacturer’s protocol. Integrity of RNA was checked on an agarose gel. Libraries were prepared using the TruSeq RNA Sample Preparation Kit (Illumina) according to the protocols recommended by the manufacturer, and each library was paired-end sequenced (2×75 bp) by using the NextSeq High Output Flow Cell - SE75 platform at the University of Missouri DNA Core. Three biological replicates were performed for each sample.

### CRC2631-PDL1 blockade treatment

To control for tumor burden, 9-12-week-old TRAMP animals were imaged and sorted into four groups (*N=12*/group). Animals were intravenously infused with PBS, 2.5×10^7^ CFU of CRC2631, or 0.5 mg Invivomab (murine anti-PDL1 antibodies; BXCELL, #BE0101) alone or in combination with 2.5×10^7^ CFU of CRC2631. Animals received one injection every three days for a total of four doses. To evaluate the effect of the therapy on tumor size, animals were MRI scanned 21 days after the completion of treatment. These MRI images were used to compare tumor sizes between groups and to determine metastatic incidences in lymph nodes and lungs.

### Canine studies

Four 13-month-old male beagles were administered one dose of 4×10^6^ CRC2631 by intravenous injection. Plasma samples were taken pre-administration (0 h) and at 2, 24, 96, and 168 h after CRC2631 administration. A small animal Maxi Panel, which covers glucose (mg/dL), urea nitrogen (mg/dL), creatinine (mg/dL), sodium (mEq/L), potassium (mEq/L), chloride (mEq/L), bicarbonate (mEq/L), anion gap (mEq/L), albumin (g/dL), total protein (g/dL), globulin (g/dL), calcium (mg/dL), phosphorus (mg/dL), cholesterol (mg/dL), total bilirubin (mg/dL), ALT (U/L), ALP (U/L), and CK (U/L), was performed and analyzed by the MU Veterinary Medical Diagnostic Laboratory (Columbia, MO).

### Genome sequencing and analysis

CRC2631 was grown in a dry shaker in 10mL of liquid LB +200 µg/ml thymine +50 µg/mL kanamycin for 24 hours at 37 °C, 220 rpm. The overnight culture was split. One volume was used for extraction of chromosomal DNA using the standard Wizard genomic DNA prep kit (Promega) protocol (CRC2631). The other volume was suspended in sterile PBS for intravenous tail injection of 2.5×10^7^ CRC2361 into groups of 11-15-week-old B6 TRAMP(+) mice that were euthanized for tissue collection at 96 or 190 h post CRC2631 injection. Following biodistribution analysis protocols, isolated colonies of CRC2631 were identified after plating prostate tumor tissue homogenates on selective LB +200 µg/ml thymine +50 µg/mL kanamycin plates for 24 h at 37 °C. These colonies were grown in 10 mL of liquid LB +200 µg/ml thymine +50 µg/mL kanamycin media for 24 h at 37 °C, 220 rpm. Four representative TRAMP prostate tumor-passaged overnight cultures from individual mice were used for extraction of chromosomal DNA as described above (CRC2631a-d). Parental and prostate tumor-passaged chromosomal genomic DNA were sequenced following the standard Next Generation Sequencing NovaSeq 2×100 protocol (Illumina) and aligned against the parental LT2 and associated stable pSLT reference sequences (NCBI: NC_003197.2, NC_003277.2) using breSeq (v0.53.1), freeBayes (v1.3.2), and TIDDIT (v2.10.0) to identify SNP mutations and structural variations present in the tumor-passaged strains (CRC2631a-d) but not in the CRC2631 injection strain. Integrative Genomics Viewer (v.2.8.0) was used to create graphical representations of SNP mutations and structural variations in all sequenced strains.

### Mathematical modeling

An estimated value of 1.75 for *λ* is interpreted as the expected number of total SNPs for CRC2631 over a 96 h interval. Next, we consider the probability of individual genes showing a SNP over a 96 h interval. It is very likely that the genes comprising CRC2631 do not all have the same probability of developing a SNP. Thus, we consider the average probability of a gene developing a SNP using our estimates for *λ*. Under our Poisson approximation, *λ*=np_avg_, where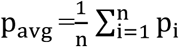 is the probability of the i^th^ gene developing a SNP, and n=4538. A *λ* estimate of 1.75 yields an estimate of 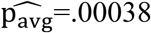 likewise, 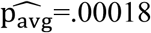 and 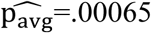 for the upper and lower bounds of the corresponding 95% credible interval for *λ*. Applying a geometric distribution to these p_avg_ estimates, we obtain an expected value of 10391 days before a SNP develops in an average gene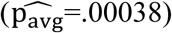, likewise 22145 days 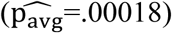 and 6141 days 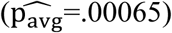.

### Cell viability assay

Prostate cells (10^4^ per well) in their respective media were seeded in 96-well plates. The cells were allowed to adhere and recover for approximately 18 h after which they were treated with 10^4^ CFU of CRC2631 for 4 h. At the completion of treatment, media were replaced with gentamycin (40 µg/mL)-containing media for 1 h to eliminate extracellular bacteria. Cells were then washed twice with 1x PBS and cell viability was measured using the MTT Cell Proliferation Assay (ATCC® 30-1010K), as per the manufacturer’s recommendation.

### Statistical analyses

All statistical analyses (student’s t test comparisons, ANOVA of mean weight differences over time) were performed using GraphPad Prism software (v6.0h).

## Abbreviations

ANOVA: analysis of variance
BCT: bacterial cancer targeting
bp: base pair
B6: C57BL/6-Tg(TRAMP)8247Ng/J mice
B6FVB: C57BL/6-Tg(TRAMP)8247Ng/J x FvBNHsd mice
CCD: charged couple device
CFU: colony forming units
CK: creatinine kinase
CSF2: colony stimulating factor 2
CXCL: chemokine ligand
Cy5: Cyanine 5
DAPI: 4′,6-diamidino-2-phenylindole
DNA: deoxyribonucleic acid
EMH: extramedullary hematopoiesis
h: hour
hpi: hours post injection
H&E: hematoxylin and eosin
IL6: interleukin 6
IL1*β*: interleukin 1 beta
INF*γ*: interferon gamma
IP: intraperitoneal
IV: intravenous
IVIS: in vivo imaging system
kV: kilovolts
LD50: median lethal dose
mEq: milliequivalents
min: minute
MRI: magnetic resonance imaging
MTD: maximum tolerated dose
NA: numerical aperture
nd: none detected
NEPC: neuroendocrine prostate cancer
nt: no treatment
OD: optical density
PALN: proper axillary lymph node
PBS: phosphate buffered saline
PD1: programmed death cell protein 1
PDL1: program death ligand-1
RFP: red fluorescent protein
RNA: ribonucleic acid
rpm: revolutions per minute
SD: standard deviation
SNP: short nucleotide polymorphism
SV40: *simian virus* 40
TIL: tumor-infiltrating lymphocytes
TNF*α*: tumor necrosis factor alpha
TRAMP: Transgenic Adenocarcinoma of Mouse Prostate
TPM: transcripts per million
U: units

## Author Contributions

YCC, RAK, and BD-M were responsible for experimental design. RAK and BD-M were responsible for carrying out all experimental work with the following exceptions: RAK and EG conducted cytokine and flow cytometry analyses, RAK, BD-M, LL, and LM conducted MRI assays, and CPD-S conducted mathematical modeling of CRC2631 risk probabilities. AAB performed NGS data analyses. RAK, BD-M, and YCC wrote the manuscript.

## Acknowledgements

We acknowledge the Yale Center for Genomic Analysis (New Haven, CT) for RNAseq and genomic sequencing, William Spollen (MU Institute for Data Science and Informatics, Columbia, MO) for assistance with SNP mapping, Melody Kroll (MU Division of Biological Sciences, Columbia MO) for manuscript proofing, and Dennis Lubahn (MU Department of Biochemistry, Columbia MO) for the gift of TRAMP breeding stock.

---

**Supplemental Table 1.**
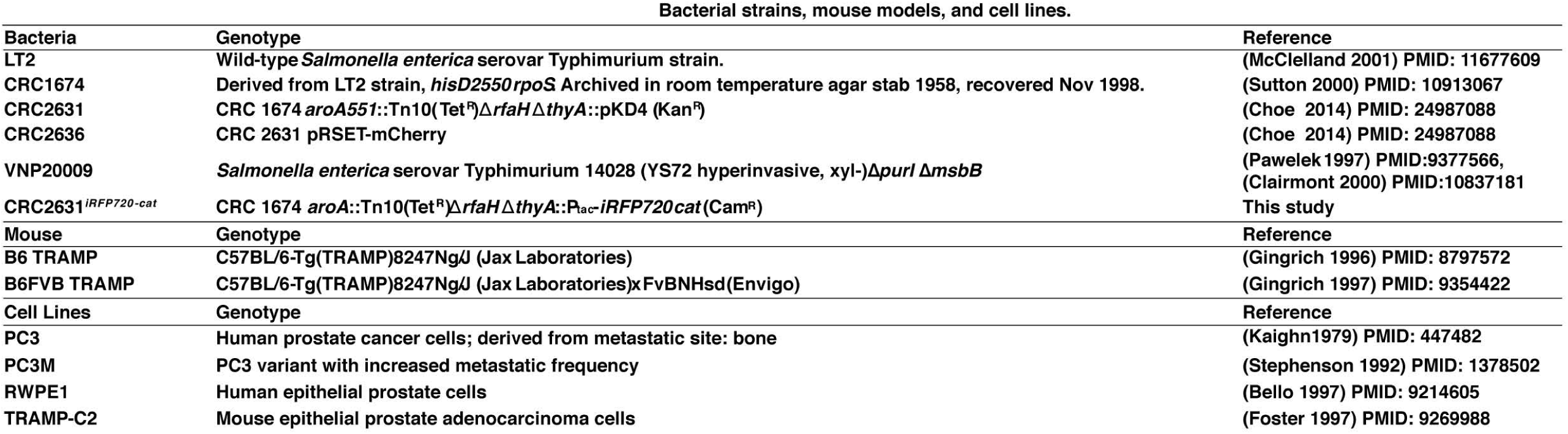
Bacterial strains, mouse models, and cell lines

**Supplemental Figure 1.**
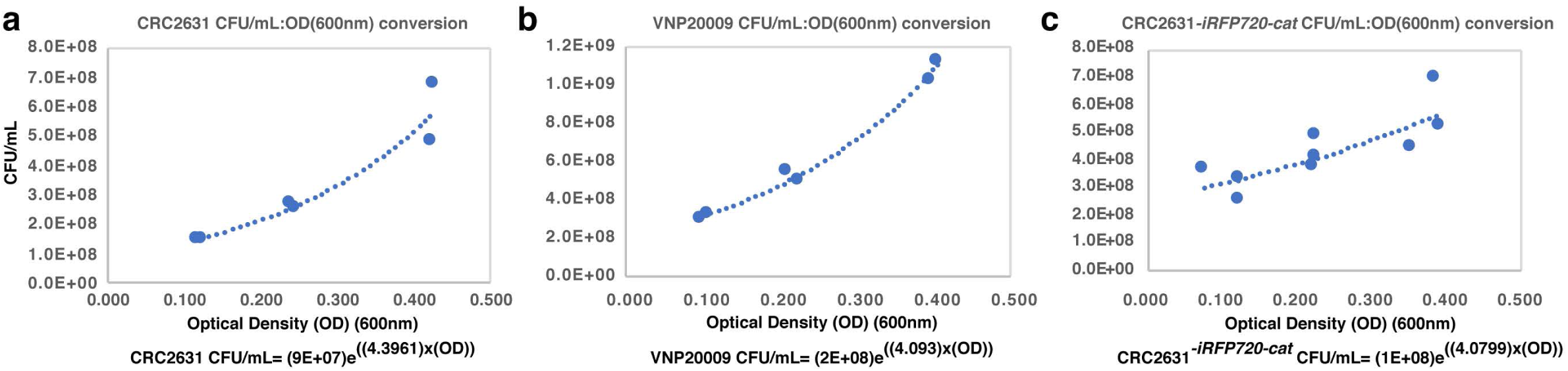
Determination of optical density conversion equations. Best fit curve equations (OD to viable cells/mL) of (a) CRC2631, (b) VNP20009, and (c) CRC2631^-*iRFP720-cat*^ independent clonal populations suspended in PBS at three different 600nm optical densities (OD) after growth for 24 h in liquid culture and viable cells/mL determined by plating dilution series of each culture on plates containing appropriate selective antibiotics (see methods) and enumerated after 30 h incubation at 37 °C.

**Supplemental Figure 2.**
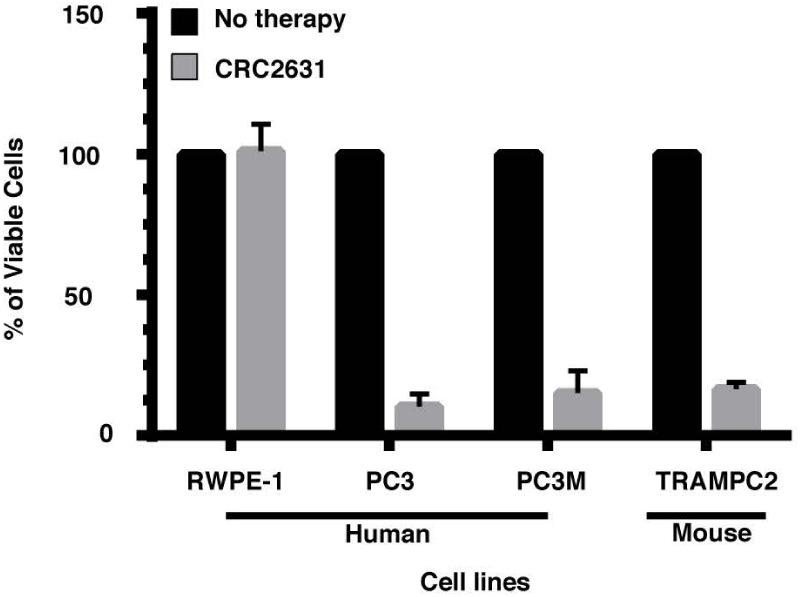
CRC2631 specifically targets human and mouse prostate cancer cells. Human benign prostate (RWPE-1), prostate cancer and murine cancer cells (10^4^) were treated with 10^4^ CFU of CRC2631 for 4 h at 37 °C and then washed. Cell viability was assessed using an MTT assay. Results represent the mean ±SD of three trials performed in triplicate.

**Supplemental Figure 3.**
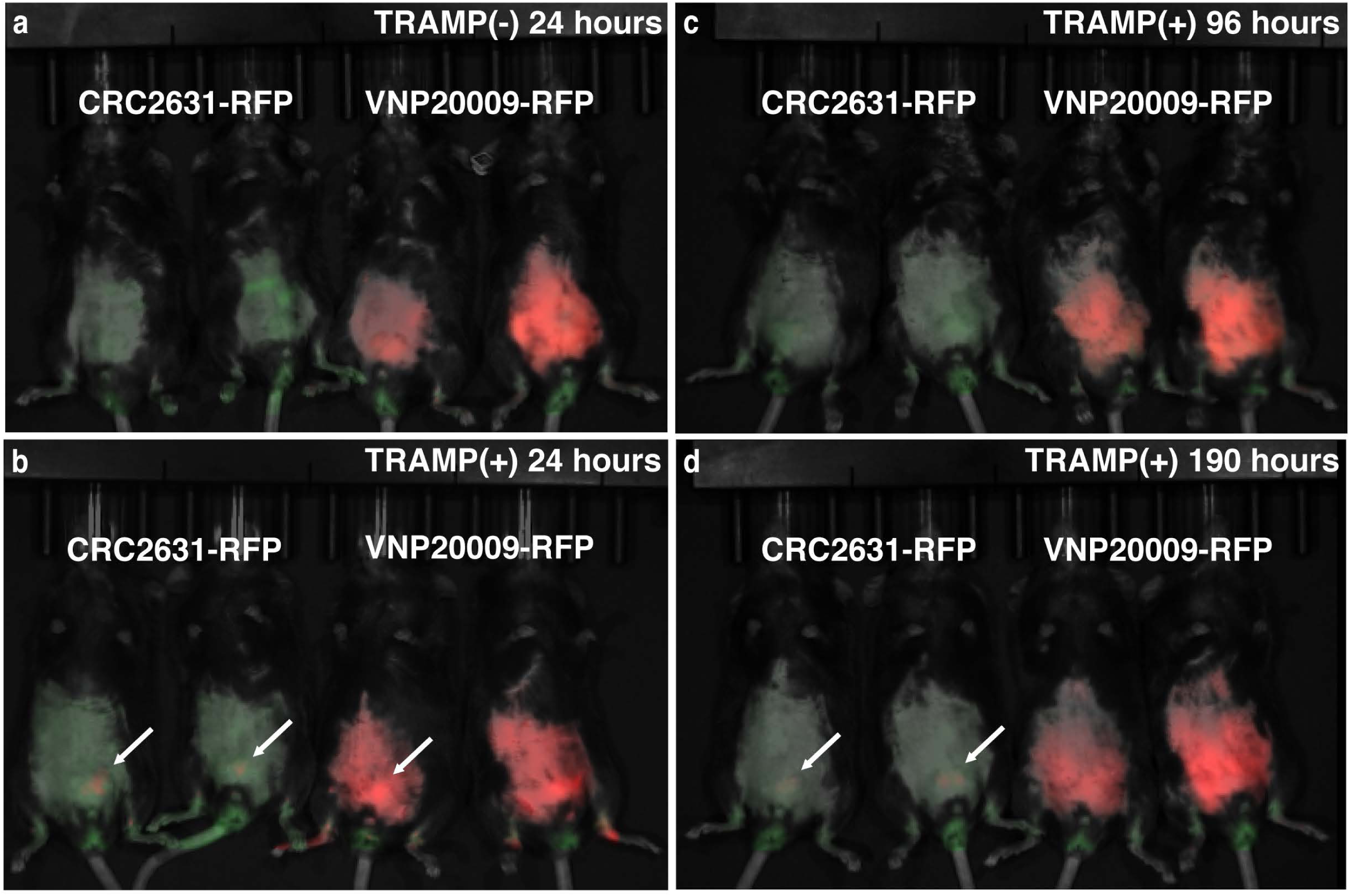
Comparative CRC2631 and VNP20009 biodistribution in B6 TRAMP animals. Qualitative localization and persistence of CRC2631 and VNP20009 expressing mCherry red fluorescent protein (RFP) in (a) B6 TRAMP(-) or (b-d) B6 TRAMP(+) mice bearing primary prostate tumors. Mice (*N=2*) were IP treated with 1 × 10^6^ CFU of CRC2631 or VNP20009 expressing mCherry red fluorescent protein (RFP). Using an IVIS *in vivo* fluorescent imaging system, living mice were scanned at (b) 24, (c) 96, and (d) 190 hours post injection to detect CRC2631 or VNP20009 associated mCherry RFP signal. Red = CRC2631 or VNP20009 associated mCherry signal. Green = tissue autofluorescence. (a) CRC2631 does not persist after 24 hours in B6 TRAMP(-) mice. (b) CRC2631 successfully colonizes the primary prostate tumor (arrows) at 24hpi in B6 TRAMP(+) mice. (c) CRC2631 mCherry signal becomes undetectable at 96hpi, but (d) re-emerges at 190hpi, demonstrating persistence in the B6 TRAMP (+) primary prostate tumor model.

**Supplemental Figure 4.**
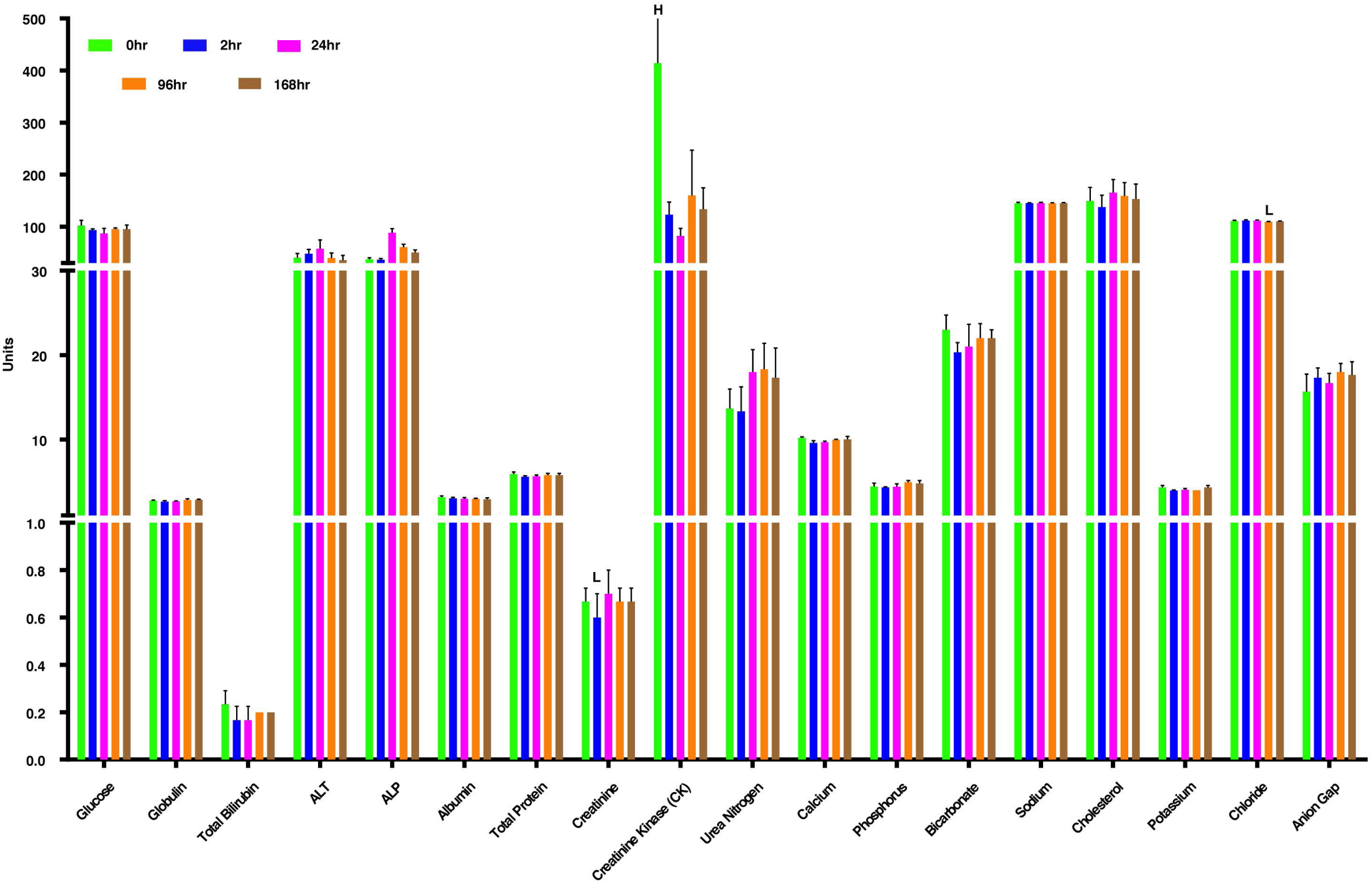
Toxicological assessment of CRC2631 in canine models. Three 13-month-old male beagles were IV administered 4×10^6^ CRC2631 and plasma samples collected at 0, 2, 24, 96, and 168 h time points. A small animal Maxi Panel was performed to evaluate pathological response to CRC2631 injection. The chart shows the mean levels of plasma chemistry components in the three dogs to identify significant pathologies in organ tissue or metabolic function. All panels that included mean results outside of calibrated normal ranges (L=Low, H=High) are shown. Mean levels of creatinine below normal range at two hours post CRC2631 injection was not significantly different from initial levels (*p*<0.374). One dog exhibited high levels of creatinine kinase (CK) before injection of CRC2631 but CK levels were within normal range from 2-168 hours post injection and mean CK level changes from pre-injection to post-injection was not significantly different at 2 h (*p*<0.372), 24 h (*p*<0.316), 96 h (*p*<0.436), or 168 h (*p*<0.389). Mean chloride levels at 96 hpi were below normal range but this was not significantly different from initial chloride levels (*p*<0.230). Chemistry panels indicate no significant pathologies in organ tissue or metabolic function as a result of IV CRC2631 injections into dogs.

## References

1. Palumbo MO, Kavan P, Miller WH, Jr., Panasci L, Assouline S, Johnson N, Cohen V, Patenaude F, Pollak M, Jagoe RT, Batist G. Systemic cancer therapy: achievements and challenges that lie ahead. Front Pharmacol. 2013; 4: 57. doi: 10.3389/fphar.2013.00057.

2. Sonpavde G, Wang CG, Galsky MD, Oh WK, Armstrong AJ. Cytotoxic chemotherapy in the contemporary management of metastatic castration-resistant prostate cancer (mCRPC). BJU Int. 2015; 116: 17–29. doi: 10.1111/bju.12867.

3. Pawelek JM, Low KB, Bermudes D. Tumor-targeted Salmonella as a novel anticancer vector. Cancer Res. 1997; 57: 4537–44. doi:

4. Clairmont C, Lee KC, Pike J, Ittensohn M, Low KB, Pawelek J, Bermudes D, Brecher SM, Margitich D, Turnier J, Li Z, Luo X, King I, et al. Biodistribution and genetic stability of the novel antitumor agent VNP20009, a genetically modified strain of Salmonella typhimurium. J Infect Dis. 2000; 181: 1996–2002. doi: 10.1086/315497.

5. Zheng LM, Luo X, Feng M, Li Z, Le T, Ittensohn M, Trailsmith M, Bermudes D, Lin SL, King IC. Tumor amplified protein expression therapy: Salmonella as a tumor-selective protein delivery vector. Oncol Res. 2000; 12: 127–35. doi:

6. Luo X, Li Z, Lin S, Le T, Ittensohn M, Bermudes D, Runyab JD, Shen SY, Chen J, King IC, Zheng LM. Antitumor effect of VNP20009, an attenuated Salmonella, in murine tumor models. Oncol Res. 2001; 12: 501–8. doi:

7. Toso JF, Gill VJ, Hwu P, Marincola FM, Restifo NP, Schwartzentruber DJ, Sherry RM, Topalian SL, Yang JC, Stock F, Freezer LJ, Morton KE, Seipp C, et al. Phase I study of the intravenous administration of attenuated Salmonella typhimurium to patients with metastatic melanoma. J Clin Oncol. 2002; 20: 142–52. doi:

8. Kazmierczak RA, Gentry B, Mumm T, Schatten H, Eisenstark A. Salmonella Bacterial Monotherapy Reduces Autochthonous Prostate Tumor Burden in the TRAMP Mouse Model. PLoS One. 2016; 11: e0160926. doi: 10.1371/journal.pone.0160926.

9. McClelland M, Sanderson KE, Spieth J, Clifton SW, Latreille P, Courtney L, Porwollik S, Ali J, Dante M, Du F, Hou S, Layman D, Leonard S, et al. Complete genome sequence of Salmonella enterica serovar Typhimurium LT2. Nature. 2001; 413: 852–6. doi:

10. Eisenstark A. Genetic diversity among offspring from archived Salmonella enterica ssp. enterica serovar typhimurium (Demerec Collection): in search of survival strategies. Annu Rev Microbiol. 2010; 64: 277–92. doi: 10.1146/annurev.micro.091208.073614.

11. Sutton A, Buencamino R, Eisenstark A. rpoS mutants in archival cultures of Salmonella enterica serovar typhimurium. J Bacteriol. 2000; 182: 4375–9. doi:

12. Edwards K, Linetsky I, Hueser C, Eisenstark A. Genetic variability among archival cultures of Salmonella typhimurium. FEMS Microbiol Lett. 2001; 199: 215–9. doi:

13. Tracy BS, Edwards KK, Eisenstark A. Carbon and nitrogen substrate utilization by archival Salmonella typhimurium LT2 cells. BMC Evol Biol. 2002; 2: 14. doi:

14. Porwollik S, Wong RM, Helm RA, Edwards KK, Calcutt M, Eisenstark A, McClelland M. DNA amplification and rearrangements in archival Salmonella enterica serovar Typhimurium LT2 cultures. J Bacteriol. 2004; 186: 1678–82. doi:

15. Hu K, Artsimovitch I. A Screen for rfaH Suppressors Reveals a Key Role for a Connector Region of Termination Factor Rho. mBio. 2017; 8. doi: 10.1128/mBio.00753-17.

16. Wang L, Jensen S, Hallman R, Reeves PR. Expression of the O antigen gene cluster is regulated by RfaH through the JUMPstart sequence. FEMS Microbiol Lett. 1998; 165: 201–6. doi: 10.1111/j.1574-6968.1998.tb13147.x.

17. Hoiseth SK, Stocker BA. Aromatic-dependent Salmonella typhimurium are non-virulent and effective as live vaccines. Nature. 1981; 291: 238–9. doi: 10.1038/291238a0.

18. Kok M, Buhlmann E, Pechere JC. Salmonella typhimurium thyA mutants fail to grow intracellularly in vitro and are attenuated in mice. Microbiology. 2001; 147: 727–33. doi:

19. Mir R, Jallu S, Singh TP. The shikimate pathway: review of amino acid sequence, function and three-dimensional structures of the enzymes. Crit Rev Microbiol. 2015; 41: 172–89. doi: 10.3109/1040841X.2013.813901.

20. Berman-Booty LD, Knudsen KE. Models of neuroendocrine prostate cancer. Endocr Relat Cancer. 2015; 22: R33–49. doi: 10.1530/ERC-14-0393.

21. Foster BA, Gingrich JR, Kwon ED, Madias C, Greenberg NM. Characterization of prostatic epithelial cell lines derived from transgenic adenocarcinoma of the mouse prostate (TRAMP) model. Cancer Res. 1997; 57: 3325–30. doi:

22. Gingrich JR, Barrios RJ, Kattan MW, Nahm HS, Finegold MJ, Greenberg NM. Androgen-independent prostate cancer progression in the TRAMP model. Cancer Res. 1997; 57: 4687–91. doi:

23. Gingrich JR, Barrios RJ, Morton RA, Boyce BF, DeMayo FJ, Finegold MJ, Angelopoulou R, Rosen JM, Greenberg NM. Metastatic prostate cancer in a transgenic mouse. Cancer Res. 1996; 56: 4096–102. doi:

24. Gingrich JR, Greenberg NM. A transgenic mouse prostate cancer model. Toxicol Pathol. 1996; 24: 502–4. doi: 10.1177/019262339602400414.

25. Greenberg NM, DeMayo FJ, Sheppard PC, Barrios R, Lebovitz R, Finegold M, Angelopoulou R, Dodd JG, Duckworth ML, Rosen JM, et al. The rat probasin gene promoter directs hormonally and developmentally regulated expression of a heterologous gene specifically to the prostate in transgenic mice. Mol Endocrinol. 1994; 8: 230–9. doi:

26. Levine AJ, Momand J, Finlay CA. The p53 tumour suppressor gene. Nature. 1991; 351: 453–6. doi: 10.1038/351453a0.

27. Aggarwal R, Zhang T, Small EJ, Armstrong AJ. Neuroendocrine prostate cancer: subtypes, biology, and clinical outcomes. J Natl Compr Canc Netw. 2014; 12: 719–26. doi: 10.6004/jnccn.2014.0073.

28. Gingrich JR, Barrios RJ, Foster BA, Greenberg NM. Pathologic progression of autochthonous prostate cancer in the TRAMP model. Prostate Cancer Prostatic Dis. 1999; 2: 70–5. doi:

29. Kaplan-Lefko PJ, Chen TM, Ittmann MM, Barrios RJ, Ayala GE, Huss WJ, Maddison LA, Foster BA, Greenberg NM. Pathobiology of autochthonous prostate cancer in a pre-clinical transgenic mouse model. Prostate. 2003; 55: 219–37. doi: 10.1002/pros.10215.

30. Maia MC, Hansen AR. A comprehensive review of immunotherapies in prostate cancer. Crit Rev Oncol Hematol. 2017; 113: 292–303. doi: 10.1016/j.critrevonc.2017.02.026.

31. Coutermarsh-Ott SL, Broadway KM, Scharf BE, Allen IC. Effect of Salmonella enterica serovar Typhimurium VNP20009 and VNP20009 with restored chemotaxis on 4T1 mouse mammary carcinoma progression. Oncotarget. 2017; 8: 33601–13. doi: 10.18632/oncotarget.16830.

32. Shcherbakova DM, Verkhusha VV. Near-infrared fluorescent proteins for multicolor in vivo imaging. Nat Methods. 2013; 10: 751–4. doi: 10.1038/nmeth.2521.

33. Sun C, Mezzadra R, Schumacher TN. Regulation and Function of the PD-L1 Checkpoint. Immunity. 2018; 48: 434–52. doi: 10.1016/j.immuni.2018.03.014.

34. Kythreotou A, Siddique A, Mauri FA, Bower M, Pinato DJ. Pd-L1. J Clin Pathol. 2018; 71: 189–94. doi: 10.1136/jclinpath-2017-204853.

35. Gao J, Ward JF, Pettaway CA, Shi LZ, Subudhi SK, Vence LM, Zhao H, Chen J, Chen H, Efstathiou E, Troncoso P, Allison JP, Logothetis CJ, et al. VISTA is an inhibitory immune checkpoint that is increased after ipilimumab therapy in patients with prostate cancer. Nat Med. 2017; 23: 551–5. doi: 10.1038/nm.4308.

36. Pasero C, Gravis G, Guerin M, Granjeaud S, Thomassin-Piana J, Rocchi P, Paciencia-Gros M, Poizat F, Bentobji M, Azario-Cheillan F, Walz J, Salem N, Brunelle S, et al. Inherent and Tumor-Driven Immune Tolerance in the Prostate Microenvironment Impairs Natural Killer Cell Antitumor Activity. Cancer Res. 2016; 76: 2153–65. doi: 10.1158/0008-5472.CAN-15-1965.

37. Gray A, de la Luz Garcia-Hernandez M, van West M, Kanodia S, Hubby B, Kast WM. Prostate cancer immunotherapy yields superior long-term survival in TRAMP mice when administered at an early stage of carcinogenesis prior to the establishment of tumor-associated immunosuppression at later stages. Vaccine. 2009; 27 Suppl 6: G52–9. doi: 10.1016/j.vaccine.2009.09.106.

38. Felgner S, Frahm M, Kocijancic D, Rohde M, Eckweiler D, Bielecka A, Bueno E, Cava F, Abraham WR, Curtiss R, 3rd, Haussler S, Erhardt M, Weiss S. aroA-Deficient Salmonella enterica Serovar Typhimurium Is More Than a Metabolically Attenuated Mutant. mBio. 2016; 7. doi: 10.1128/mBio.01220-16.

39. Hadrup S, Donia M, Thor Straten P. Effector CD4 and CD8 T cells and their role in the tumor microenvironment. Cancer Microenviron. 2013; 6: 123–33. doi: 10.1007/s12307-012-0127-6.

40. Han S, Toker A, Liu ZQ, Ohashi PS. Turning the Tide Against Regulatory T Cells. Front Oncol. 2019; 9: 279. doi: 10.3389/fonc.2019.00279.

41. Kumar P, Bhattacharya P, Prabhakar BS. A comprehensive review on the role of co-signaling receptors and Treg homeostasis in autoimmunity and tumor immunity. J Autoimmun. 2018; 95: 77–99. doi: 10.1016/j.jaut.2018.08.007.

42. Sambi M, Bagheri L, Szewczuk MR. Current Challenges in Cancer Immunotherapy: Multimodal Approaches to Improve Efficacy and Patient Response Rates. J Oncol. 2019; 2019: 4508794. doi: 10.1155/2019/4508794.

43. Van Dessel N, Swofford CA, Forbes NS. Potent and tumor specific: arming bacteria with therapeutic proteins. Ther Deliv. 2015; 6: 385–99. doi: 10.4155/tde.14.113.

44. Kasinskas RW, Forbes NS. Salmonella typhimurium specifically chemotax and proliferate in heterogeneous tumor tissue in vitro. Biotechnol Bioeng. 2006; 94: 710–21. doi: 10.1002/bit.20883.

45. Chandrasekaran EV, Xue J, Neelamegham S, Matta KL. The pattern of glycosyl- and sulfotransferase activities in cancer cell lines: a predictor of individual cancer-associated distinct carbohydrate structures for the structural identification of signature glycans. Carbohydr Res. 2006; 341: 983–94. doi: 10.1016/j.carres.2006.02.017.

46. Datsenko KA, Wanner BL. One-step inactivation of chromosomal genes in Escherichia coli K-12 using PCR products. Proc Natl Acad Sci U S A. 2000; 97: 6640–5. doi:

47. Edwards RA, Keller LH, Schifferli DM. Improved allelic exchange vectors and their use to analyze 987P fimbria gene expression. Gene. 1998; 207: 149–57. doi: 10.1016/s0378-1119(97)00619-7.

48. Greenberg NM, DeMayo F, Finegold MJ, Medina D, Tilley WD, Aspinall JO, Cunha GR, Donjacour AA, Matusik RJ, Rosen JM. Prostate cancer in a transgenic mouse. Proc Natl Acad Sci U S A. 1995; 92: 3439–43. doi: 10.1073/pnas.92.8.3439.

49. Inoue Y, Izawa K, Kiryu S, Tojo A, Ohtomo K. Diet and abdominal autofluorescence detected by in vivo fluorescence imaging of living mice. Mol Imaging. 2008; 7: 21–7. doi:

50. Hedges AJ. Estimating the precision of serial dilutions and viable bacterial counts. Int J Food Microbiol. 2002; 76: 207–14. doi: 10.1016/s0168-1605(02)00022-3.

